# *Wolbachia* as agents of extensive mtDNA lineage sharing between species through multiple infection

**DOI:** 10.1101/2024.04.03.587946

**Authors:** Víctor Noguerales, Brent C. Emerson

## Abstract

*Wolbachia* can manipulate arthropod host reproduction, triggering the homogenisation of mtDNA variation within species and introgression between hybridising species through indirect selection. While fixation within species of mtDNA variants linked to *Wolbachia* infections has been documented, a broader understanding of the potential consequences of *Wolbachia* infection through hybridisation is limited. Here we evaluate *Wolbachia* transmission through hybridisation as a mechanistic explanation for extensive mtDNA paraphyly between two species of iron-clad beetle (Zopheridae). Our analyses reveal a complex pattern of mitochondrial variation, supporting the introgression of at least five mtDNA lineages from *Tarphius canariensis* into *T. simplex*, in a background of a shared *Wolbachia* infection across both species. Genetic clustering and demographic simulations reveal a clear pattern of nuclear differentiation between species, a limited signature of historical gene flow, and the eastwards range expansion of *T. simplex* across the existing distribution of *T. canariensis.* These results are consistent with hybridisation during early stages of secondary contact, during which *Wolbachia* infection facilitated recurrent mtDNA introgression events. These results highlight the complex restructuring of mitochondrial differentiation across invertebrate species that can result from bacterial endosymbiotic infections, a phenomena with potentially profound impacts for the disciplines of phylogeography and species delimitation.

## INTRODUCTION

The sharing of genetic variation among closely related species is an expected outcome of either recent speciation and incomplete lineage sorting (ILS), or hybridisation, or a combination of both (Toews & Brelsford 2012; Good *et al*. 2015). In addition to these two processes, indirect selection on mitochondrial DNA (mtDNA) arising from linkage disequilibrium with inherited microorganisms also has the potential to homogenise patterns of genetic variation between species (Cariou *et al*. 2017). Strong indirect selection for mtDNA haplotypes can emerge from the manipulation of host reproduction towards the survival of the daughters of females infected with parasitic symbionts. The most common form of host reproductive manipulation is cytoplasmic incompatibility (Hurst & Jiggins 2005; Kiefer *et al*. 2022), which may be either uni- or bidirectional (Engelstädter & Telschow 2009; Wang *et al*. 2022; Hochstrasser 2023). Such mating incompatibilities arise between individuals with and without cytoplasmic endosymbiotic parasites, whereby matings between uninfected females and infected males are incompatible, while matings between infected females and uninfected or similarly infected males are compatible (Bordenstein *et al*. 2001; Jiggins 2003). Under these conditions, infected females are reproductively favoured, and mtDNA variants associated with infected females hitchhike with symbionts that undergo selective sweeps within populations. The homogenising effects of such infections for mtDNA are well understood, where mtDNA variation within a species is replaced by a single haplotype associated with the initial infection (*e.g.*, Turelli *et al*. 1992; Raychoudhury *et al*. 2010). This will ultimately lead to fixation of the symbiont associated haplotype within the host species, if the symbiont has sufficient drive to spread, and host populations are sufficiently connected by dispersal, and infection duration is sufficient for complete spread of the infection to occur. While early speculation suggested that infection turnover might be rapid (Hurst & Jiggins 2005), more recent evidence points to infection durations that may extend over evolutionary time-scales (Bailly-Bechet *et al*. 2017). This is supported by reports of high levels of mtDNA divergence within some infected species (*e.g.,* Hinojosa *et al*. 2019, 2022), indicative of infection persisting beyond the fixation time for the initial symbiont-associated haplotype, and thus co-occurring with haplotype variation that has arisen through *de novo* mutations from the haplotype associated with the original sweep.

Estimates of the proportion of arthropod species infected by *Wolbachia* largely fall within a range of 40-50% (Zug & Hammerstein 2012; Weinert *et al*. 2015; Lefoulon *et al*. 2016; Bailly-Bechet *et al*. 2017), but could be higher than 60% when taking into account differing infection frequencies and sampling effects (Hilgenboecker *et al*. 2008). While transmission of *Wolbachia* within species is vertical, transmission between species, and hence novel infection, is horizontal, either through predation of infected individuals, parasitism, shared ecological niches or hybridisation (Kaur *et al*. 2021). Among these pathways, hybridisation is likely to be more efficient, as it directly involves the reproductive machinery used in vertical transmission, and may be frequent, depending upon the extent of reproductive isolation among species.

Transfer and fixation of mtDNA between species through *Wolbachia* infection is now reasonably well understood. The closely related butterfly species *Acraea encedana* and *A. encedon* are both infected by *Wolbachia.* The species are distinct based on morphology and nuclear genetic variation, but *A. encedana* and *A. encedon* individuals with the same *Wolbachia* infection have identical mtDNA, while uninfected *A. encedon* individuals have a distinct mtDNA genome (Jiggins 2003). This sharing of mtDNA genomes in a background of nuclear genomic segregation can be explained by rare hybridisation events followed by indirect selection for a single mtDNA haplotype via *Wolbachia* (Hurst & Jiggins 2005). Even if F_1_ progeny are of low fitness, any successful backcrossing with an uninfected parental species may open the door to the spread of the mtDNA from the infected parental species through symbiont drive and hitchhiking (Bech *et al*. 2021). *Wolbachia*-associated transfer and fixation of mtDNA between species has also been observed in butterflies of the genera *Iphiclides* (Gaunet *et al*. 2019), *Lycaeides* (Gompert *et al*. 2008) and *Polytremis* (Jiang *et al*. 2018), *Diplazon* parasitoid wasps (Klopfstein *et al*. 2016), *Altica* leaf beetles (Jäckel *et al*. 2013) and *Drosophila*, with introgression between *D. simulans* and *D. mauritania* (Rousset & Solignac 1995; Ballard 2000).

The relationship between *Wolbachia* infection and indirect selective sweeps of mtDNA is also well recognised (*e.g*., Raychoudhury *et al*. 2009; Cariou *et al*. 2017; Dincă *et al*. 2019; Martin *et al*. 2020), and has been suggested to be a regular event in insects (Hurst & Jiggins 2005). Multiple independent *Wolbachia* infections of arthropod species have also been reported (*e.g.*, Werren *et al*. 1995; Reuter & Keller 2003; Narita *et al*. 2007; Miyata *et al*. 2020), although these are thought to be less common than single infections, due to their lower stability leading to limited persistence time (Engelstädter & Telschow 2009). However, in their analysis of swallowtail butterflies, Gaunet *et al*. (2019) document a sequential infection by two *Wolbachia* strains from *Iphiclides podalirius* to *I. feisthamelii*, such that mtDNA variation in *I. feisthamelii*, derived from a historical sweep is now being replaced by an ongoing sweep associated with the second infection.

*Wolbachia* may facilitate mtDNA introgression between closely related species through hybridisation (*e.g.*, Rousset & Solignac 1995; Jiggins 2003; Narita *et al*. 2006; Charlat *et al*. 2009; Dyer *et al*. 2011; Jäckel *et al*. 2013), and the results of Gaunet *et al*. (2019) and Miyata *et al*. (2020) highlight how hybridisation frequency and spatial factors can lead to complex patterns of mtDNA relatedness among species involving multiple shared mtDNA lineages. Given recent evidence that *Wolbachia* infection through hybridisation can drive more complex patterns of mtDNA relatedness (Gaunet *et al*. 2019; Miyata *et al*. 2020) than classically observed single lineage sweeps (Rousset & Solignac 1995; Ballard 2000; Jiggins 2003; Hurst & Jiggins 2005), there is a need to better understand *Wolbachia*-mediated mtDNA dynamics between closely related species.

Here we seek to further our understanding by focusing on two closely related and morphologically distinct species of iron-clad beetle (Zopheridae: Colydiinae) from the genus *Tarphius* that occur in sympatry within laurel cloud forest on the Canary island of Tenerife. Phylogenetic analysis of mtDNA variation for *T. canariensis* and *T. simplex* has revealed that individuals from neither species segregate as a monophyletic group, with a complex pattern of mtDNA paraphyly across both species that has previously been suggested to be the result of either ILS or hybridisation (Emerson *et al*. 2000; Salces-Castellano *et al*. 2020). Here we expand the sampling of Salces-Castellano *et al*. (2020) for both species for a total 108 *T. canariensis* and 81 *T. simplex* from 19 sites, for which 16 sites present both species in sympatry (Fig. 1). We generate mtDNA sequences and ddRAD-seq data for all individuals to test hypotheses of: (i) hybridisation between both species, and (ii) ILS using demographically-derived predictions. We also interrogate the nuclear data for the presence of *Wolbachia* sequences across all individuals. We then apply demographic modelling and predictions from population genetic theory to the nuclear data to provide a spatiotemporal framework for the establishment of species sympatry. Finally, the minimum number of mtDNA introgression events between *T. canariensis* and *T. simplex* is estimated from a network analysis of mtDNA sequences.

**Figure 1.**
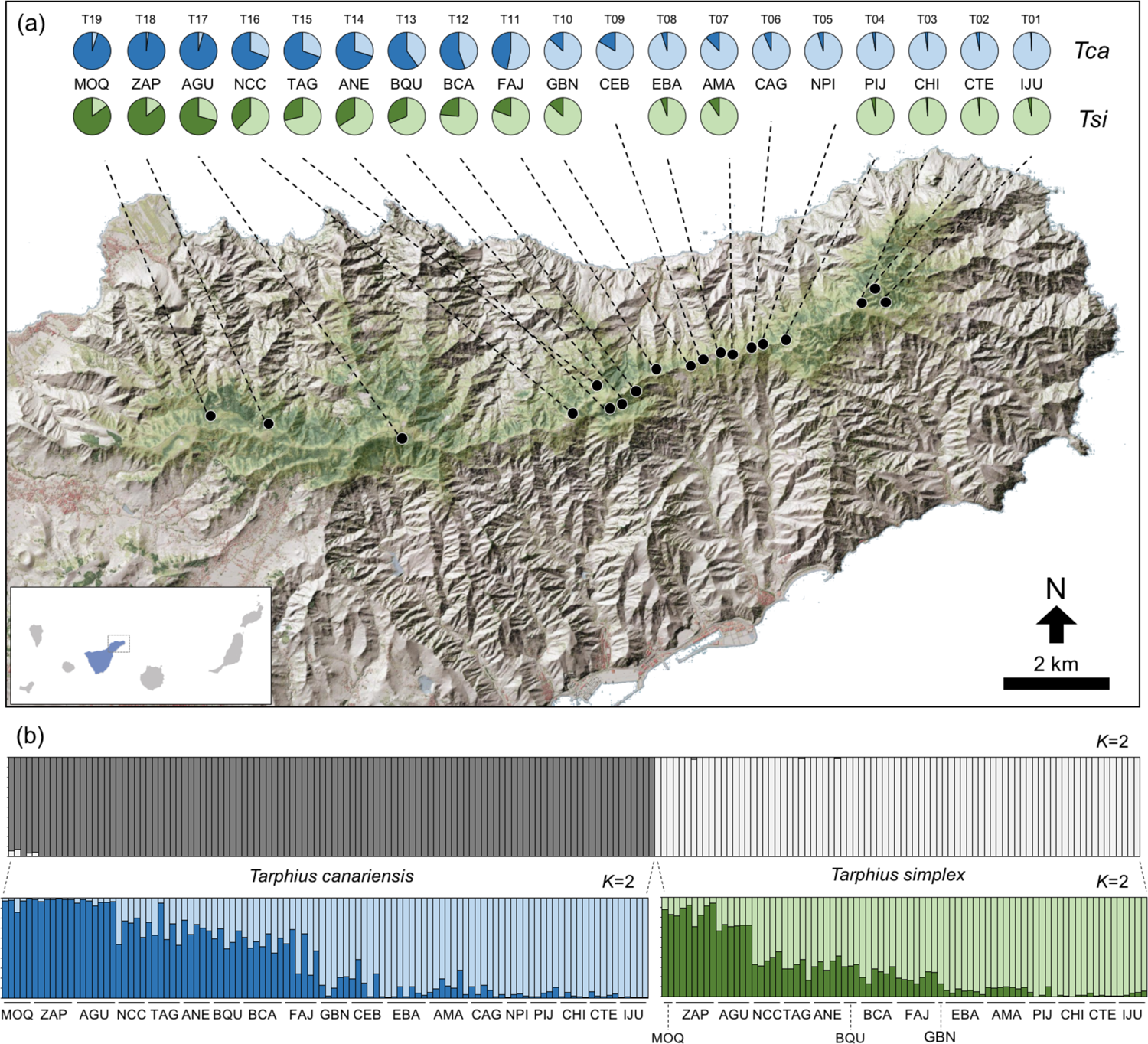
Panel (a) shows the geographical location of sampling sites within the Anaga peninsula of Tenerife, and population ancestry coefficients (pie charts) for *Tarphius canariensis* (*Tca*) and *T. simplex* (*Tsi*) as inferred in STRUCTURE, assuming two ancestral populations (*K* = 2) within each species. Missing pie charts for *Tsi* represent sampling sites where the species was not found. Inset map shows the Canary Islands and the location of the Anaga peninsula within Tenerife (in blue). Panel (b) shows the hierarchical genetic clustering results. The upper bar plot depicts the ancestry coefficients per individual when both species are analysed together, with lower bar plots describing inferences when each species is analysed independently assuming *K* = 2. Thin vertical black lines separate individuals, which are partitioned into *K*-colored segments representing the probability of belonging to a given cluster. Population codes as in Table S1.

We find a pattern of complete mitochondrial introgression involving at least five mtDNA lineages from *T. canariensis* to *T. simplex.* This has occurred in a background of strong nuclear differentiation between both species, with a model supporting only limited historical gene flow in the absence of contemporary admixture signatures, but with a shared *Wolbachia* infection across both species. Demographic and population genetic analyses reveal *T. simplex* to have more recently established within the Anaga peninsula, within which dispersal-limited range expansion would have led to a period of incremental contact with populations of *T. canariensis* across its existing range within the peninsula. Our results provide insight on how *Wolbachia* infection and species-specific demographic histories can jointly interact to drive complex patterns of mitochondrial relatedness that might not intuitively be associated with *Wolbachia*. We discuss the broader implications of our results, which suggest that pathogenic symbionts may play a greater role in explaining shared patterns of mtDNA variation among species than might otherwise be thought.

## MATERIAL AND METHODS

### Sample collection

Specimens of *Tarphius canariensis* and *T. simplex* were sampled from 19 sites along the dorsal ridge of the Anaga peninsula (Fig. 1), yielding a total of 108 *T. canariensis* and 81 *T. simplex*, with individuals of both species sampled together at 16 sites (Table S1). See Supplementary Methods S1 for further details on sampling.

### Mitochondrial and ddRADseq data

Genomic DNA was extracted from each individual using the Biosprint DNA Blood Kit (Qiagen) on a Thermo KingFisher Flex automated extraction instrument. The barcode region of the mitochondrial DNA (mtDNA) cytochrome c oxidase subunit I (COI) was amplified using the primers *Fol-degen-for* and *Fol-degen-rev* (Yu *et al*. 2012). PCR conditions are described in Table S2. PCR products were purified with enzymes *ExoI* and *rSAP* (New England Biolabs, Ipswich, MA, USA), Sanger sequenced (Macrogen, Madrid, Spain), edited with GENEIOUS PRIME 2021.1.1 and aligned using MAFFT (FFT-NS-i method; Katoh & Standley 2013).

A double-digestion restriction-site associated DNA sequencing (ddRADseq) protocol, as described Salces-Castellano *et al*. (2021), was applied. In brief, individual DNA extracts were digested with the restriction enzymes *MseI* and *EcoRI* (New England Biolabs), genomic libraries were pooled at equimolar ratios, size selected for fragments between 200-300 base pairs (bp), and then paired-end sequenced (150 bp) on an Illumina NovaSeq6000 (Novogene, Cambridge, UK).

### Haplotype networks using mtDNA data

Mitochondrial DNA sequences were collapsed into haplotypes using FABOX 1.61 (Villesen 2007) and a haplotype network was constructed using POPART 1.7 (Leigh & Bryant 2015) with statistical parsimony, as described in Clement *et al*. (2000). The haplotype network was rooted using an individual of *T. canariensis* from the neighbouring island of La Palma (Emerson *et al*. 2000). Species-haplotype associations were mapped onto the rooted haplotype network to jointly infer haplotype ancestry and species association, together with the minimum number of mtDNA haplotypes that are, or have been, shared between both species. Under a model of indirect selection for mtDNA haplotypes by parasitic symbiont transmission between species, the direction of introgression between *T. canariensis* and *T. simplex* was inferred from the order of species state change from the root of the network. This is analogous to the approach taken by García-Olivares *et al*. (2017) to infer the minimum number and direction of dispersal events for mtDNA lineages between islands (see Figure 3 in García-Olivares *et al*. 2017).

### Bioinformatic analyses for ddRADseq data

Raw sequences were demultiplexed, quality filtered and *de novo* assembled using IPYRAD 0.9.81 (Eaton & Overcast 2020). Detailed information on sequence assembly and data filtering is provided in Supplementary Methods S2. Unless otherwise indicated all downstream analyses were performed with a clustering threshold of sequence similarity of 0.85 (*clust_threshold*), discarding loci that were not present in at least 70% of individuals (*min_samples_locus*), and unlinked SNPs (*i.e.*, a single SNP per locus).

### Genomic clustering and phylogenomic inference

Population genomic structure was inferred using SNP data and the Bayesian Markov Chain Monte Carlo (MCMC) clustering method implemented in the program STRUCTURE 2.3.3 (Pritchard *et al*. 2000), assuming correlated allele frequencies and a model of admixture. A hierarchical approach was applied to identify the underlying genomic variation at the within-species level, as performed in Ortego *et al*. (2021). First, a global analysis was undertaken to test for signatures of hybridisation between both species across all sampling sites, and then species were analysed individually. For both levels of analysis, log probabilities of Pr(X|*K*) (Pritchard *et al*. 2000) and Δ*K* (Evanno *et al*. 2005) statistics were used to infer the number of ancestral populations (*K*), as recommended by Gilbert *et al*. (2012) and Janes *et al*. (2017). Supplementary Methods S3 provides further details on genomic clustering analyses. In addition, we visualised the major axis of genomic variation both between and within species with a Principal Component Analysis (PCA) using the ‘*gl.pcoa’* function as implemented in the package *dartR* (Mijangos *et al*. 2022) and the R version 4.2.2 (R Core Team 2022).

Phylogenetic relationships among the ancestral populations inferred with STRUCTURE were reconstructed using the coalescent-based method for species tree estimation implemented in SNAPP (Bryant *et al*. 2012). We excluded individuals of admixed ancestry by restricting analyses to individuals with a high probability of assignment (*q*-value > 85%) to an ancestral population. Supplementary Methods S3 provides additional details on phylogenomic inference.

### Population genetic differentiation

For sampling sites with ≥4 individuals (Table S1), pairwise genomic differentiation was measured among all such sites with the *F*_ST_ and *N*_ST_ estimators for ddRADseq and mtDNA data, respectively. *F*_ST_ and *N*_ST_ values were calculated with the ‘*gl.fst.pop*’ function of the R package *dartR*, and DNASP 5.10.1 (Librado & Rozas 2009), respectively. Statistical significance of estimators was tested with 1000 bootstrapping replicates.

Correlation between *F*_ST_ and *N*_ST_ matrices, and their respective correlations with geographic distance (isolation-by-distance scenario, IBD), were evaluated using Mantel tests implemented in the R package *vegan* (Oksasen *et al*. 2022). Geographic distances between all sampling sites were calculated with the geodesic method implemented in the R package *geodist* (Padgham 2021).

### Analyses of genomic diversity

For sampling sites with ≥4 individuals (Table S1), ddRADseq data was used to estimate genomic diversity with the unbiased expected heterozygosity (u*H*_E_) and nucleotide diversity (π) estimators. u*H*_E_ was estimated with the ‘*gl.report.heterozygosity*’ function of the R package *dartR*, whereas π was estimated in DNASP 6.12.03 (Rozas *et al*. 2017) using as input the .*allele* file generated with IPYRAD.

Geographical clines of genomic diversity (Guo 2012) were tested for by contrasting measures of population genetic variability (u*H*_E_ and π) and spatial variables (longitude and latitude), using generalised linear models (GLMs) in R software. Models were fitted with the weighted least square method to weight each observation proportional to its sample size (*e.g*., Noguerales *et al*. 2018).

### Testing alternative demographic models

Coalescent demographic modelling was used to statistically test for the fit of data to alternative scenarios of divergence and migration. Four demes corresponding to the co-distributed West and East ancestral populations of each species were used, and models were constructed to estimate the timing of divergence and the magnitude of gene flow among demes (Fig. S1). Analyses were conducted using the same subset of individuals and demes used for phylogenomic inference in SNAPP (see Supplementary Methods S3). On the basis of the SNAPP topology, scenarios of divergence in strict isolation (Model A) and alternative models considering both intraspecific (Model B), and ancestral interspecific migration (Model C), were evaluated. Models assuming contemporary interspecific gene flow were constructed involving migration: (i) only between the West demes (Model D), (ii) between West and East demes, separately (Model E), and (iii) among West and East demes (Model F) (Fig. S1). These migration models considered either symmetrical or asymmetrical migration matrices between the two taxa. The composite likelihood of the observed data was estimated, given a specified model using the site frequency spectrum (SFS) and the simulation-based approach implemented in FASTSIMCOAL 2.5.2.21 (Excoffier *et al*. 2013). Details on composite likelihood estimation, model selection approach and calculation of confidence intervals for parameter estimates under the most-supported model are described in Supplementary Methods S4.

### Analyses of *Wolbachia* infection

The raw ddRADseq data was interrogated for *Wolbachia* sequences using CENTRIFUGE 1.0.4 (Kim *et al*. 2016). The combination of this approach with ddRADseq data from insects has been proven to extract reliable information on the infection of host individual samples by bacterial endosymbionts (Hinojosa *et al*. 2022, 2023). Briefly, reads matching to *Wolbachia* genomes were extracted and subjected to a comprehensive quality filtering pipeline as detailed in Supplementary Methods S5. Potential *Wolbachia* sequences were further verified with the *blastn* function from BLAST+ 2.15.0 (Camacho *et al*. 2009; *e.g.*, Lucek *et al*. 2020), and only those unambiguously identified as *Wolbachia* were *de novo* assembled and curated using GENEIOUS. Supplementary Methods S5 provides detailed information on analyses for identifying *Wolbachia* sequences and subsequent data filtering and curation.

## RESULTS

### Mitochondrial and ddRADseq data

A total of 185 mtDNA sequences were obtained, corresponding to 483 bp within the COI region, which resolved to 95 unique haplotypes. Maximum uncorrected genetic divergences within species (2.3% and 2.7% for *T. canariensis* and *T. simplex*) were similar to that observed between species (2.7%). Illumina sequencing provided a total of 379.54 M reads across all 189 individuals, with an average of 1.78 M reads per sample (SD = 1.00 M) (Fig. S2). After the different filtering and assembly steps, each individual retained an average 34,685 clusters (SD = 13.06), with a mean depth per locus of 23.50 (SD = 8.01) across individuals.

### MtDNA haplotype network

The rooted COI haplotype network revealed a minimum of five mtDNA lineages shared between *T. canariensis* and *T. simplex*. Under a model of introgression, all five lineages are inferred to have been transferred from *T. canariensis* to *T. simplex* (Fig. 2). While haplotypes of *T. canariensis* segregate into two lineages that are, with limited exceptions, phylogeographically consistent with the West and East regions within Anaga (*sensu* Salces-Castellano *et al*. 2020), such phylogeographic structure was not reflected within the mtDNA variation of *T. simplex* (Fig. 2). This lack of phylogeographic structure for *T. simplex* was accompanied by strong signatures for *T. simplex* haplotypes being derived from western *T. canariensis* haplotypes for four out of the five shared lineages, with the ancestry of the fifth lineage being consistent with either a western or eastern origin from *T. canariensis* (Fig. 2).

**Figure 2.**
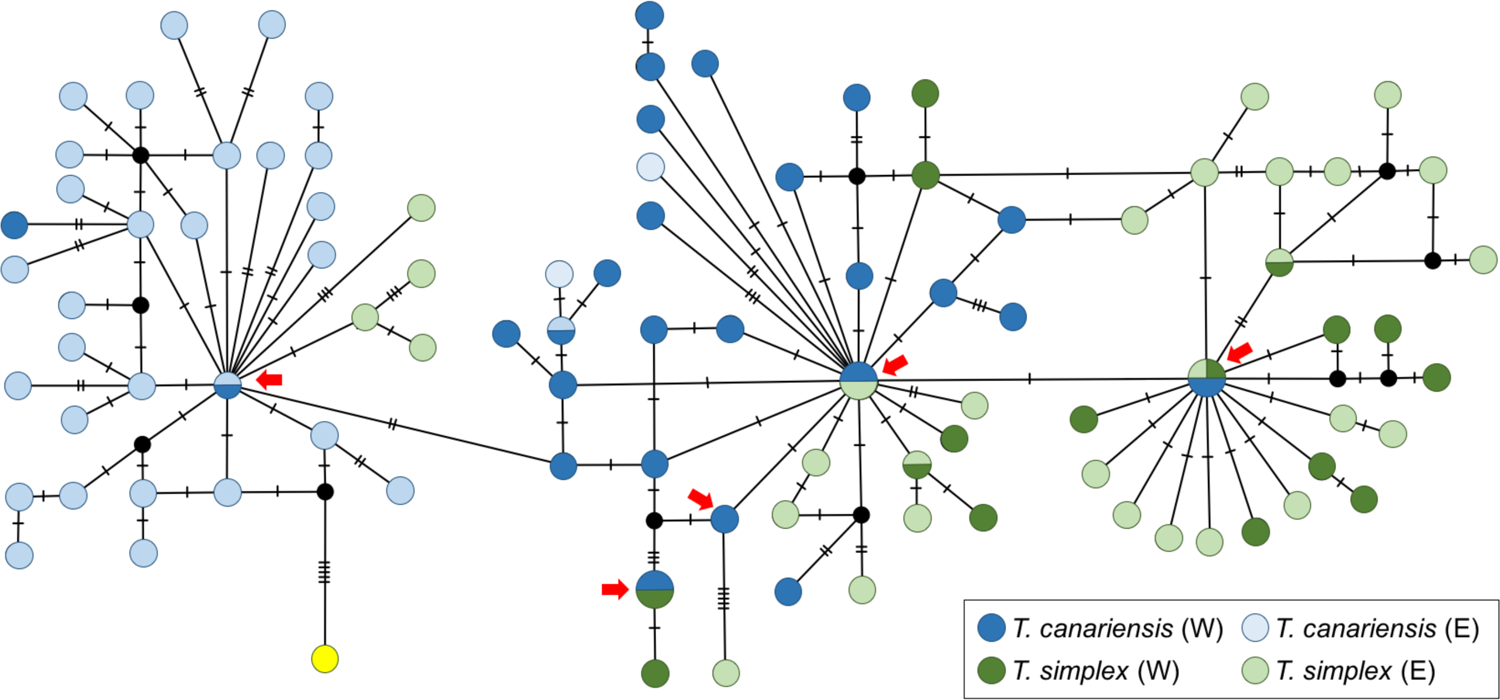
Haplotype network depicting relationships between mtDNA COI haplotypes sampled from both *Tarphius canariensis* and *T. simplex*. Dark blue and dark green represent haplotypes of *T. canariensis* and *T. simplex*, respectively, sampled in the West (W) region. Lighter coloured-circles depict haplotypes of each respective species that were sampled in the East (E) region. Red arrows show either shared haplotypes between the two species or independent events representing *T. simplex* haplotypes derived from *T. canariensis*. Outgroup is shown in yellow. West (W) and East (E) regions were defined according to STRUCTURE inferences for *T. canariensis* (Fig. 1), with the delimiting point between regions falling on the sampling site of FAJ (Table S1).

**Figure 3.**
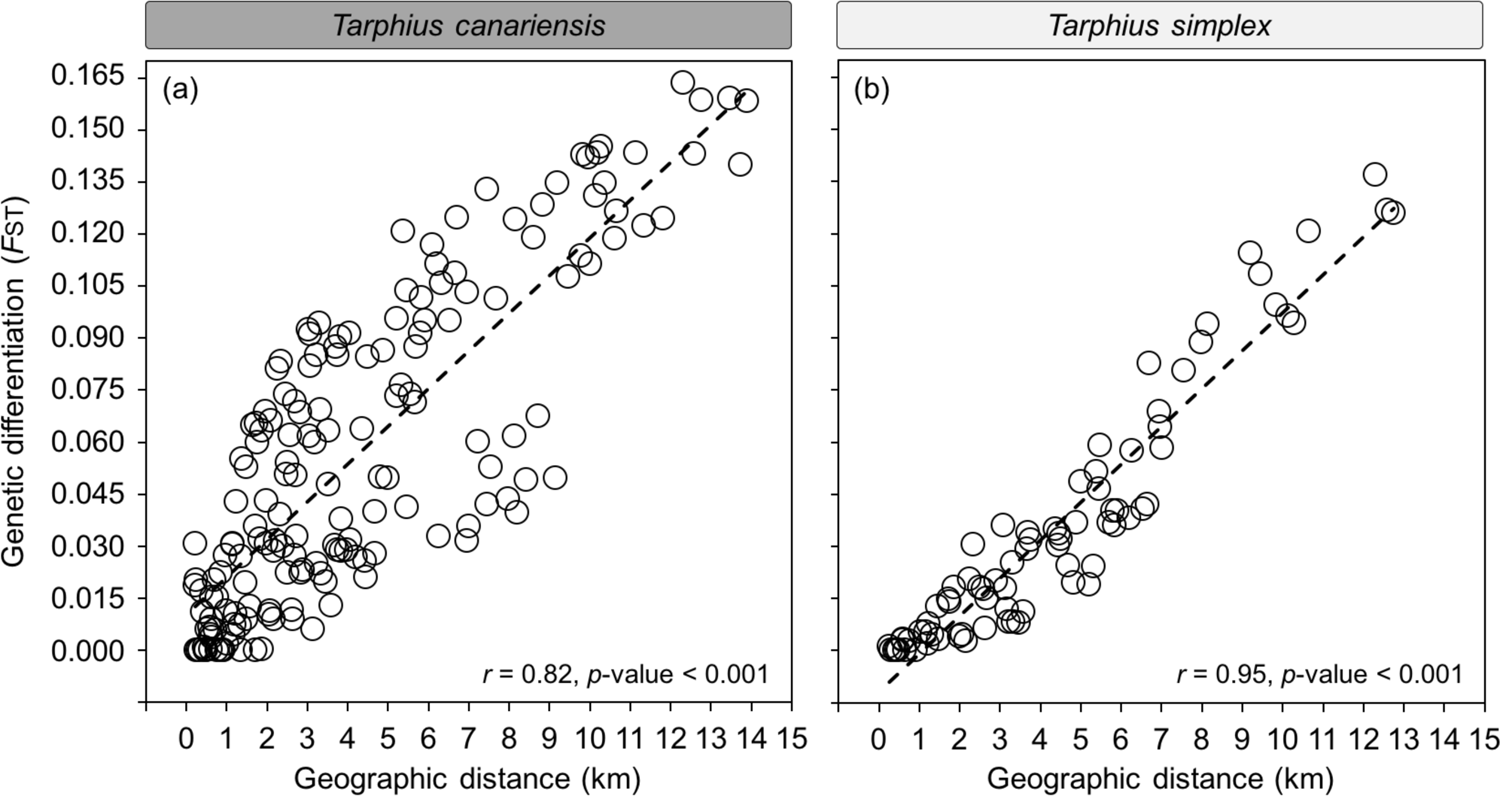
Relationship between genetic differentiation (*F*_ST_) of nuclear genomic variation and geographic distance between populations for each of the two species, *T. canariensis* (panel a) and *T. simplex* (panel b).

### Genomic clustering and phylogenomic inference

Analyses across all individuals of both species with STRUCTURE identified the most likely number of ancestral populations to be two, according to log probability and Δ*K* statistics (Fig. S3), with each taxonomic species representing an ancestral population with all individuals presenting high single ancestry coefficients (Fig. 1), in line with patterns of individual similarity observed with PCA (Fig. S4). While STRUCTURE analyses provided consistent results of species cohesiveness and limited hybridisation, 4 individuals of *T. canariensis* (MOQ) and 2 individuals of *T. simplex* (ZAP, TAG) presented signatures of admixed ancestry (1.4 < *q*-value < 7.3%; Fig. 1), suggesting they may be representative of historical introgression events within the western region of Anaga.

Further STRUCTURE analyses at the intraspecific level revealed genomic variation to be hierarchically organised, with two inferred ancestral populations within each species (Fig. S3), corresponding to western and eastern sampling sites, with a cline of co-ancestry between them (Fig. 1). While this broad pattern of single ancestry and admixture is shared between the two species, *T. simplex* presents a more gradual gradation from East to West, with individuals of high single ancestry assignment found only in the East (Fig. 1). In comparison, *T. canariensis* presents multiple sampling sites characterised by high single ancestry assignment in both the East and the West, with a sharp geographic transition of ancestry assignment coinciding with the sampling site of FAJ (Fig. 1). Consistent with inferences from STRUCTURE, PCAs revealed genomic variation within each species to be structured into West and East genetic groups (Fig. S4), with less pronounced differentiation observed in *T. simplex*. Differences in allele frequencies are described along the PC1, on which individuals from central sites showed an intermediate position in concordance with the species-specific gradients of admixture detected in STRUCTURE (Fig. 1, Fig. S4).

After excluding individuals of admixed ancestry (*q*-values < 0.85), and grouping individuals by ancestry, four genetic groups of similar sample size (*T. canariensis*: West, n = 12, East, n = 10; *T. simplex*: West, n = 9, East, n = 10) were obtained. These groups were composed of conspecific individuals from the neighbouring sampling sites MOQ and ZAP (West), and IJU and CTE (East), and were used for phylogenomic inference and demographic modelling analyses (Supplementary Methods S3). Phylogenomic analyses in SNAPP supported the monophyly of each species and yielded a relative shallower divergence between the West and East groups of *T. simplex* that that observed in *T. canariensis* (Fig. S5). Further analyses in TREESETANALYZER showed that the 95% credible set of trees was represented by this single topology. Different gamma prior distributions yielded similar topologies and relative branch lengths. Estimates of population size (θ) from SNAPP were markedly lower in both groups of *T. simplex* (0.069, 0.077) compared to *T. canariensis* (0.136, 0.122).

### Population genetic differentiation and genetic diversity

Genetic differentiation (*F*_ST_) estimates between sampling sites ranged from 0.0 to 0.164 in *T. canariensis*, and from 0.0 to 0.137 in *T. simplex*. Analyses of mtDNA variation revealed *N*_ST_ estimates ranging from 0.0 to 0.660 in *T. canariensis*, and from 0.0 to 0.425 in *T. simplex. F*_ST_ values were strongly and significantly correlated with geographic distances in both species (Fig. 3), consistent with an important role of isolation-by-distance (IBD). This was particularly striking in the case of *T. simplex* (*r* = 0.95, *p* < 0.001), with isolation by distance explaining less of the observed variation within *T. canariensis* (*r* = 0.82, *p* < 0.001). *N*_ST_ values were only significantly correlated with geographic distances in *T. canariensis* (*r* = 0.55, *p* < 0.001; Fig. S7), with no support for an isolation by distance relationship for mtDNA variation within *T. simplex* (*r* = 0.27, *p* = 0.76). Accordingly, *F*_ST_ and *N*_ST_ estimates were only significantly correlated in *T. canariensis* (*r* = 0.76, *p* < 0.001, Fig. S6). Together these results suggest that geographic distance is a strong predictor of relatedness between populations for both species, but that this relationship is disrupted for the mitochondrial genome in *T. simplex*. Further analyses including sampling sites at the distribution margins of both species revealed that *N*_ST_ values were lower between species within West and East regions, respectively, than *N*_ST_ values within species across their ranges, inconsistent with an scenario of incomplete lineage sorting (ILS) (Fig. S7).

Nucleotide diversity (π) decreased significantly with longitude and latitude in both species (*p*-values < 0.028; Fig. S8), consistent with the progressive erosion of genetic variation during easternward range expansions. While this spatial pattern was consistent across both species, only *T. simplex* presented a significant geographic gradient of unbiased expected heterozygosity (*uH*_E_, Fig. S9). This is consistent with a more recent range expansion, compared to *T. canariensis,* such that allelic richness in eastern sampling sites of *T. simplex* have yet to return to equilibrium levels. Spatial patterns of genetic diversity across nuclear and mitochondrial genomes are disassociated within *T. simplex*, but are associated within *T. canariensis.* A pattern of lower nucleotide diversity (π) in the East, compared to the West, is also observed for mtDNA in *T. canariensis* (Fig. S7), but not in *T. simplex*.

### Demographic model testing

The most supported model identified with FASTSIMCOAL incorporated both intraspecific gene flow between West and East demes and symmetric interspecific gene flow within western and eastern regions (Model E1) (Fig. 4, Table S3). On the basis of a 1-year generation time, estimates from the most supported model suggest that both species diverged from a common ancestor approximately 490 Kya (95% confidence interval: 360-620 Kya). Subsequent divergence between West and East groups of *T. canariensis* is inferred to have occurred prior (200 Kya, CI: 170-250 Kya) to regional divergence within *T. simplex* (120 Kya, 95% confidence interval: 100-160 Kya), consistent with patterns of divergence inferred with SNAPP.

**Figure 4.**
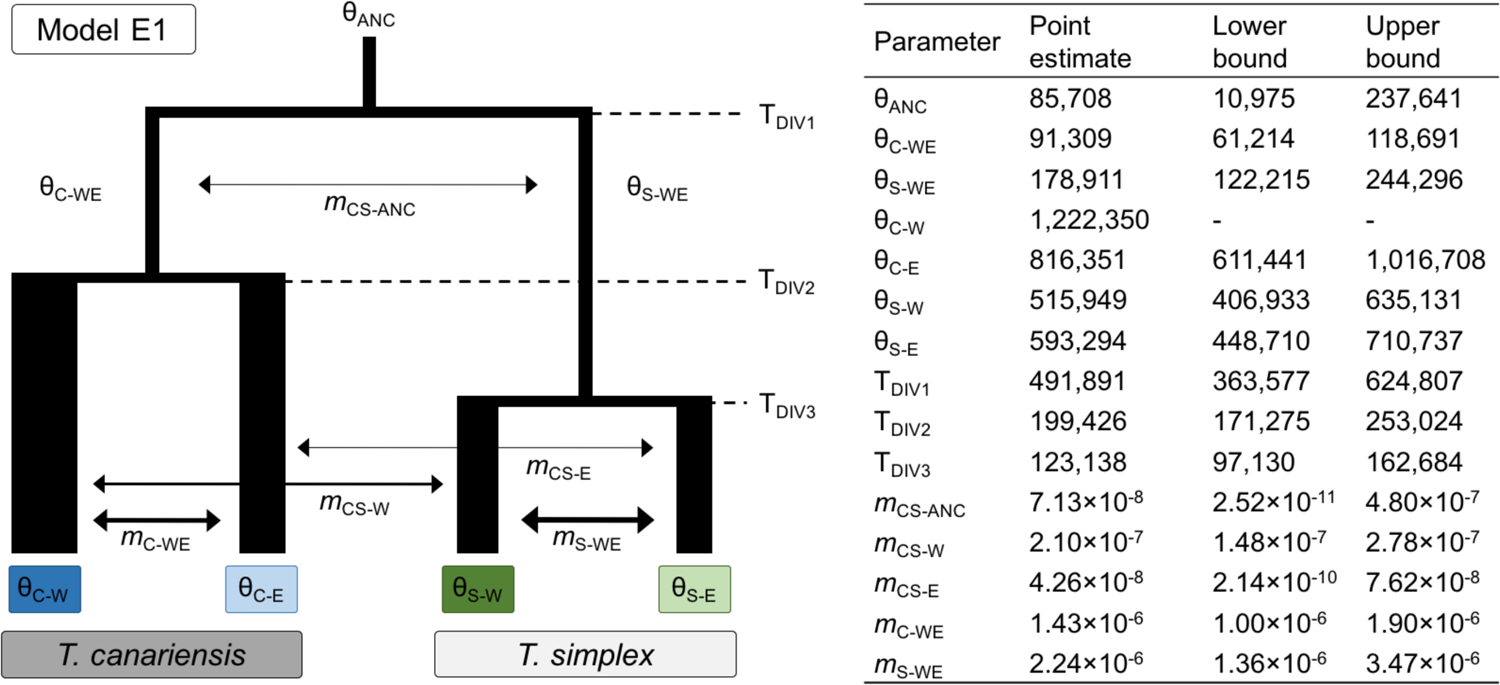
Parameters inferred from coalescent simulations with FASTSIMCOAL under the best-supported demographic model. For each parameter, its point estimate and lower and upper 95% confidence intervals are shown. Model parameters include ancestral (θ_ANC_, θ_C-WE_, θ_S-WE_) and contemporary (θ_C-E_, θ_S-W_, θ_S-E_) effective population sizes, timing of divergence (T_DIV1_, T_DIV2_, T_DIV3_) and migration rates per generation (*m*) within and between species. Width and length of branches represent θ and divergence time, respectively, with arrow thickness representing the magnitude of gene flow. Note that θ for the West genetic group (θ_C-W_) of *T. canariensis* was fixed in FASTSIMCOAL analyses to enable the estimation of other parameters.

The estimated number of migrants per generation (calculated by multiplying migration rate by population size) is low between species, but are inferred to be higher within the West region (0.13 migrants per generation) than in the East (0.02 migrants per generation), suggesting increased barriers to gene flow between the species as *T. simplex* expanded its range eastward into already established populations of *T. canariensis*. At the intraspecific level, the number of migrants per generation between West and East demes were estimated to be largely similar for *T. simplex* (0.58 migrants per generation) and *T. canariensis* (0.87 migrants per generation), with lower estimates of contemporary population size (*N*_E_) in *T. simplex* compared to *T. canariensis* (Fig. 4), in accordance with estimates from SNAPP.

### Analyses of *Wolbachia* infection

A total of 10,614 reads that were assigned to *Wolbachia* were retrieved in CENTRIFUGE, of which 10,229 were further taxonomically verified in BLAST+ (Table S4). *Wolbachia* was detected in 25 individuals, corresponding to 8 *T. canariensis* (∼7.3% prevalence) and 17 *T. simplex* (∼21.0% prevalence) with infected individuals being broadly distributed across the ranges of both species (Table S4). The number of *Wolbachia* reads per individual was not correlated with the total number of host raw reads (Spearman’s rank correlation: *r* = 0.22, *p*-value = 0.270). After sequence assembly, a total of 383 *Wolbachia* loci were obtained, of which 372 (∼97.1%) presented no variation across individuals. Of these invariant loci, 125 (∼32.6%) were sampled in both *T. canariensis* and *T. simplex.* Only 11 loci (∼2.9%) showed genetic variation, with four of them presenting variation among individuals within the same species, and the remaining seven loci presenting variation among individuals from both species (Table S4). Four of these seven loci were variant due to differences found in a single individual of *T. simplex* (siT15TAG06, Table S5), with variation represented by only one variant site in three of these four loci. The number of *Wolbachia* reads and loci recovered were positively correlated (Spearman’s rank correlation: *r*_s_ = 0.70, *p*-value < 0.001) and varied greatly across individuals, with 5 individuals from both species sampled exclusively for over 95% of all shared loci (Table S4), likely representing individuals with a higher level of infection.

## DISCUSSION

The nuclear genomes of both *Tarphius canariensis* and *T. simplex* provide for a detailed understanding of the origin of their sympatric distributions within the Anaga peninsula of Tenerife, and the history of gene flow between them. Multiple lines of evidence argue for both species ranges having extended from the western limits of the dorsal ridge of the Anaga peninsula, across to the east, with an older origin for *T. canariensis* within the peninsula, followed by a more recent establishment of *T. simplex,* resulting in their sympatry. This demographic and evolutionary context provided the opportunity for hybridisation events during the early stages of secondary contact, leading to the *Wolbachia*-mediated transfer of mtDNA lineages from *T. canariensis* to *T. simplex*. Below we discuss the contrasting patterns of mito-nuclear discordance between both species as a result of the mtDNA replacement resulting from a dynamics of introgression and *Wolbachia* transmission.

### Community assembly dynamics and sympatry

Nucleotide diversity within each species shows significant decay from their western range limits within the peninsula, across the northeastern axis of the peninsula to their eastern range limits (Fig. S8). Such decay is consistent with the loss of allelic, and thus nucleotide variation, as a consequence of range expansion (Hewitt 2004; Excoffier *et al*. 2009), and is expected to persist until mutation-drift equilibrium is restored through new allelic variation derived from *de novo* mutation. While both species are characterised by a gradient of decreasing nucleotide diversity from west to east, only *T. simplex* presents a corresponding gradient of decreasing expected heterozygosity (Fig. S9), signifying that not only has nucleotide diversity not recovered to levels seen in western populations, but allelic diversity as well (*e.g.*, Zhao *et al*. 2020).

Taken together, the above-mentioned results are consistent with a history where *T. canariensis* became established within the peninsula prior to *T. simplex*. Diminished nucleotide diversity for *T. canariensis* from west to east indicates that the timing of range expansion across the peninsula has been sufficiently recent such that eastern populations have yet to arrive at a mutation-drift equilibrium that characterises western populations. Additionally, the absence of significant differences in expected heterozygosity across the range of *T. canariensis* is consistent with sufficient time having passed since range expansion for allelic variation to have reached levels that characterise western populations (Nei *et al*. 1975; Chakraborty & Nei 1977; Austerlitz *et al*. 1997). In summary, mutational time has been sufficient to restore allelic diversity in *T. canariensis*, but insufficient to achieve allelic divergences that characterise western populations. In contrast, significantly diminished levels from west to east for both nucleotide diversity and expected heterozygosity in *T. simplex* indicate a more recent range expansion, subsequent to which there has been insufficient time for the recovery of neither nucleotide nor allelic diversity.

Phylogenomic and demographic modelling results provide further support for *T. simplex* having expanded its range into areas already populated by *T. canariensis*. Both Bayesian phylogenetic analysis and coalescent modelling recover regional divergence within *T. canariensis* that predates that of *T. simplex* (Fig. 4, Fig. S5). The geographically coincident regional divergences of both species are characteristic of many co-occurring species of Coleoptera within the cloud forest habitat of the peninsula (Salces-Castellano *et al*. 2020), and is explained by the fragmentation of suitable habit, provoked during periods of glacial climate (Salces-Castellano *et al*. 2021). Regional divergence time estimates (Fig. 4) within *T. canariensis* (≈ 200 kya) and *T. simplex* (≈ 125 kya) broadly coincide with the penultimate and ultimate interglacial periods, respectively (Jouzel *et al*. 2007; Berger *et al*. 2016), consistent with range expansions during periods of higher humid forest connectivity.

### Contrasting patterns of genomic concordance between *T. canariensis* and *T. simplex*

Both species of *Tarphius* present a strong pattern of isolation by geographic distance for nuclear genomic variation (Fig. 3), with less variation explained within *T. canariensis*, likely in part explained by its more pronounced west-east regional structure compared to *T. simplex* (Fig. 1, Fig. 4). Structuring of mtDNA variation within *T. canariensis* is broadly concordant with nuclear genomic differentiation, with mtDNA variation being significantly related to both geographic distance and nuclear genomic differentiation (Fig. S6). In contrast, mtDNA variation in *T. simplex* presents a low and non-significant relationship to both geographic distance and nuclear genomic differentiation (Fig. S6). This strong discordance between patterns of nuclear and mitochondrial genomic relatedness are indicative of independent drivers of their differentiation within *T. simplex*.

Analyses of nuclear genomic variation provide a clear picture of reproductive isolation between both species (Fig. 1, Fig. S4), involving a history of very limited gene flow (Fig. 4). In contrast to high diagnosability of species based on nuclear genomic variation, both species share mtDNA variation (Fig. 2), with higher mitochondrial differentiation among populations within each species than between both species from the same sampling sites (Fig. S7). Such striking discordance between both genomes is inconsistent with incomplete lineage sorting (ILS). First, due to the lower effective population size for the mtDNA genome compared to the nuclear genome, and thus a stronger influence of genetic drift, mtDNA ILS should be accompanied by similar, if not higher ILS, across both nuclear genomes. This is not the case, as evidenced by the single topology retrieved in the 95% tree set inferred in SNAPP (Fig. S5). Second, mtDNA haplotype variation in *T. simplex* increases from west to east along the peninsula (Fig. 2, Fig. S7), which is opposite to expectations under a model of ILS, within a scenario of eastward range expansion.

### The dynamics of mtDNA replacement

Transfer of mtDNA from *T. canariensis* to *T. simplex* is supported by two pieces of independent evidence. First, mtDNA introgression is expected to disrupt any shared patterns of mito-nuclear relatedness within the receiving species, which could be both *T. canariensis* and *T. simplex* for bidirectional introgression, of only one of the species, in the case if unidirectional introgression. As described above, *T. canariensis* presents strong relationships of relatedness for both genomes with geographic distance, and a strong correlation of mitochondrial and nuclear relatedness among populations. In contrast, while nuclear genomic relatedness is strongly correlated with geographic distance in *T. simplex*, there is no relationship of the mitochondrial genome with either, arguing for transfer of mtDNA variation from *T. canariensis* to *T. simplex.* Second, outgroup rooting of the haplotype network reveals ancestral sequences to be uniquely present within *T. canariensis*. Haplotypes sampled from *T. simplex* form a minimum of five lineages that are independently derived from haplotypes either shared with, or exclusive to *T. canariensis*.

Evidence for *Wolbachia* infection was found across both species of *Tarphius*, with infection detected in approximately 7% of *T. canariensis*, and approximately 21% of *T. simplex* (Table S4). Infection levels appear to vary among individuals, as indicated by correlated variation for the number of reads and loci assigned to *Wolbachia* (Spearman’s rank correlation: *r*_s_ = 0.70, *p*-value < 0.001), neither of which were correlated with the total number of host reads (Spearman’s rank correlations: *r*_s_ < 0.23, *p*-values > 0.270), or the average depth of cover of host loci (Spearman’s rank correlations: *r*_s_ < 0.13, *p*-values > 0.506), within individuals. Five individuals presented high infection levels, with 6 or more loci recovered, while the remaining 20 individuals (80%) were characterised by only one or two loci, suggesting that many individuals are likely to have infection levels below the detection limits of our data (Table S5).

Of the 383 *Wolbachia* loci recovered, 125 were sampled in both *T. canariensis* and *T. simplex*, with no differences between species, consistent with a shared infection. A causal relationship between this *Wolbachia* infection and mtDNA introgressions from *T. canariensis* to *T. simplex* is supported by mounting evidence for *Wolbachia*-mediated mtDNA introgression through hybridisation (*e.g.,* Rousset & Solignac 1995; Jiggins 2003; Narita *et al*. 2006; Charlat *et al*. 2009; Dyer *et al*. 2011; Jäckel *et al*. 2013; Gaunet *et al*. 2019; Miyata *et al*. 2020), together with high persistence times for *Wolbachia* infections within species (Bailly-Bechet *et al*. 2017; Hinojosa *et al*. 2019, 2022). Within this scenario, mtDNA mutational variation within *T. simplex* is a combination of existing divergence among haplotypes that were introgressed from *T. canariensis,* together with subsequent mutations within these that postdate introgression. An alternative explanation of historical introgression of mtDNA by direct selection from *T. canariensis* to *T. simplex*, followed by more recent, but independent, *Wolbachia* infection of each species would give rise to greater mtDNA homogeneity within, compared to between species, which is not observed.

### Hybridisation as a gateway for recurrent *Wolbachia* infection and mitochondrial introgression

Mitochondrial introgression mediated by *Wolbachia* transmission between hybridising species has classically been observed as single haplotype introgression (*e.g.,* Rousset & Solignac 1995; Ballard 2000; Jiggins 2003; Hurst & Jiggins 2005). In several cases, sharing of two mtDNA lineages has been reported (Gaunet *et al*. 2019; Miyata *et al*. 2020), where each lineage is associated with infection by a different strain of *Wolbachia*. In the case of Gaunet *et al*. (2019), polymorphism is transient due to sequential, and thus competing infections of different strains that induce cytoplasmic incompatibility (CI). Only in the case of Miyata *et al*. (2020) is the polymorphism likely to be stable, due the potential for coexistence of phenotypically different *Wolbachia* stains (CI and feminising) associated with the each introgressed mtDNA lineage (Dedeine *et al*. 2004; Narita *et al*. 2007; Engelstädter *et al*. 2008; Richardson *et al*. 2016). Our results highlight how stable coexistence of multiple introgressed mtDNA lineages can emerge from single strain infections. Patterns of mtDNA lineage sharing between species, such as those observed between *T. canariensis* and *T. simplex*, may be expected within a history where: (i) hybridisation was not uncommon, and; (ii) mtDNA variation was present within the *Wolbachia*-infected donor species prior to hybridisation. Within this scenario, hybridisation must sufficiently post-date infection of the first species such that variation within the first species has recovered through mutation, as summarised in Figure 5.

**Figure 5.**
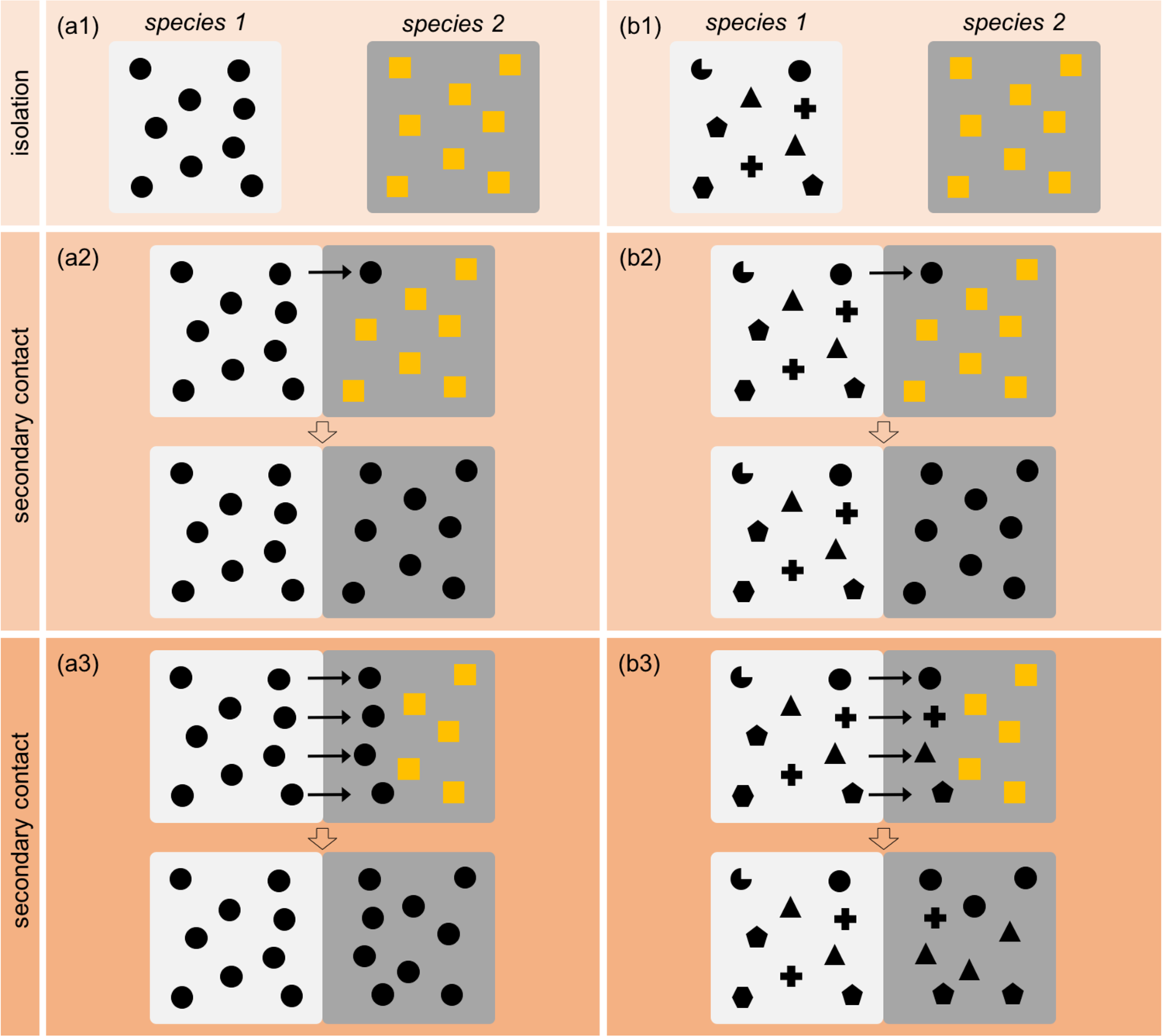
Schematic representation of differing expectations for the number of introgressed mtDNA lineages arising through *Wolbachia* infection, with regard to hybridisation frequency and mtDNA polymorphism. Species ranges are represented by white and grey boxes, within a dynamic of isolation (1) followed by secondary contact (2-3). Different shapes represent different haplotypes. *Wolbachia* infected individuals are represented by black filled shapes, while uninfected individuals are represented by orange filled shapes. Panel (a1) represents a scenario where individuals from species 1 share a unique mtDNA haplotype. Under conditions of both low (a2) and high (a3) hybridisation frequency, introgressed variation into species 2 will be low. Panel (b1) depicts a scenario where individuals from species 1 present mtDNA variation. Under this scenario, low hybridisation frequency (b2) favours the introgression of a limited number of variant haplotypes from species 1, while higher hybridisation frequency (b3) favours the introgression of higher haplotype variation from species 1.

The results presented here further our understanding of how cytoplasmic endosymbiotic parasitic infections can influence patterns of mtDNA relatedness among species. Such infections can plausibly explain complex patterns of mtDNA paraphyly that are frequently observed among invertebrate species (*e.g.,* Funk & Omland 2003; Zakharov *et al*. 2009; Gómez-Zurita *et al*. 2012; Ross 2014; Mutanen *et al*. 2016; Bilton *et al*. 2017), which may otherwise be ascribed to direct selection on the mitochondrial genome, or neutral processes of genetic drift.

## CONCLUSIONS

MtDNA has been widely used for arthropod phylogeography and species delimitation over the last four decades, and continues to play a fundamental role in characterising arthropod biodiversity through large barcoding initiatives (*e.g.*, Hendrich *et al*. 2015; Hawlitschek *et al*. 2017). While the confounding influence of *Wolbachia* for patterns of mtDNA relatedness has long been recognised (Galtier *et al*. 2009), results presented here highlight a greater challenge than might otherwise be assumed. Our study describes how hybridisation dynamics can interact with cytoplasmic endosymbiotic bacterial infection to drive complex patterns of mtDNA lineage sharing and paraphyly between what are effectively robust biological species. Approximately 40-60% of all arthropod species are estimated to be infected by cytoplasmic endosymbiotic bacteria such a *Wolbachia* (Hilgenboecker *et al*. 2008; Zug & Hammerstein 2012; Weinert *et al*. 2015; Lefoulon *et al*. 2016; Bailly-Bechert *et al*. 2017) and approximately 10% of animal species are estimated to hybridise with related taxa (Mallet 2007). We suggest that these estimates provide ample potential for the evolution of complex patterns of shared mtDNA variation among closely related arthropod species, such as those described here.

## AUTHOR CONTRIBUTIONS

BCE conceived the original idea. BCE and VN designed the research and led the study. VN analysed the data. BCE and VN wrote the manuscript.

## ACKNOWLEDGEMENTS

We thank *Centro de Supercomputación de Galicia* (CESGA) and Teide High-Performance Computing facility (TeideHPC) provided by the *Instituto Tecnológico y de Energías Renovables* (ITER), S.A. for access to computer resources. Fieldwork was supported by the *Cabildo of Tenerife* (*Expte: AFF17/23, N° Sigma: 2023-00133*). This work was supported by the Ministry of Economy and Competitiveness (MINECO) through grants CGL2013-42589-P, CGL2017-85718-P and PID2020-116788GB-I00, co-financed by FEDER. VN was supported by a *Juan de la Cierva-Formación* postdoctoral fellowship (FJC2018-035611-I) funded by MCIN/AEI/10.13039/501100011033.

## SUPPLEMENTARY MATERIAL

### SUPPLEMENTARY METHODS

#### Methods S1. Sample collection

We collected specimens of *Tarphius canariensis* and *T. simplex* species from 19 sampling sites within the laurel forest of the Anaga Peninsula, sited at the Canary island of Tenerife (Fig. 1). Geographic distance between sampling sites ranged from 0.2 km to 14 km along the dorsal ridge of the Anaga peninsula, with a maximum elevational difference between sites of 200 m (Table S1). Sampling for both species was enhanced to augment the sampling sites included in Salces-Castellano *et al*. (2020). The additional sampling effort gave rise to a total number of individuals of 189 (108 *T. canariensis,* 81 *T. simplex*), with presence of both species for 16 out of 19 sites (Table S1). Sampling was performed as described in Salces-Castellano *et al*. (2020), with minor modifications of Emerson *et al*. (2017). Specimens were preserved in 100% ethanol, taxonomically identified in the lab, and stored at –20°C till DNA extraction. Sampling was undertaken with permission of ‘Cabildo de Tenerife’ (*Expte: AFF17/23, N° Sigma: 2023-00133*).

#### Methods S2. Genomic data filtering and sequence assembly

We firstly used FASTQC 0.11.7 (Andrews 2010) to quality check raw reads. Then, raw sequences were demultiplexed, quality filtered and *de novo* assembled using IPYRAD 0.9.81 (Eaton and Overcast 2020). Only reads with unambiguous barcodes were retained (*max_barcode_mismatch*) and a stricter filter was applied to remove Illumina adapter contamination (*filter_adapters*). After trimming restriction overhangs for enzymes *EcoR1* and *MseI* (*restriction_overhang*), we converted base calls with a Phred score <20 into ambiguous sites (Ns) and discarded reads with >5 Ns (*max_low_qual_bases*). Afterwards, we clustered the retained reads within- and across samples considering a threshold of sequence similarity of 85% (*clust_threshold*) and discarded those clusters with a minimum coverage depth of less than 5 (*mindepth_majrule*) and a maximum coverage depth of more than 10,000 (*maxdepth*). Statistical base calling was performed at a minimum depth of 6 (*mindepth_statistical)*. Resulting loci shorter than 35 base pairs (bp) (*filter_min_trim_len*), containing ≥1 heterozygous sites across more than 50% individuals (*max_shared_Hs_locus*) and showing more than 20% polymorphic sites (*max_SNPs_locus*) were discarded. In a final filtering step, we only retained loci that were present in at least 70% of the samples (*min_samples_locus*), which yielded a total of 2,357, 3,161 and 7,930 unlinked SNPs, when including both *Tarphius canariensis* and *T. simplex,* only *T. canariensis* and only *T. simplex*, respectively. On average missing data in each SNP matrix was approximately 18%. Estimates of sequencing error rates and heterozygosity across individuals were on average 0.0013 (SD = 0.0004) and 0.0175 (SD = 0.0036), respectively, thus indicating an adequate specification of the clustering threshold (*clust_threshold*) parameter (Eaton & Overcast 2020).

#### Methods S3. Genomic clustering and phylogenomic inference

Population genetic structure was inferred with the Bayesian Markov Chain Monte Carlo (MCMC) clustering method implemented in the program STRUCTURE 2.3.3 (Pritchard *et al*. 2000). We ran STRUCTURE with 200,000 MCMC cycles after a burn-in step of 100,000 iterations, assuming correlated allele frequencies and admixture (Pritchard *et al*. 2000) and performing 30 independent runs for each value of *K* ancestral populations (from *K* = 1 to *K* = 5). The most likely number of ancestral populations was estimated after retaining the 10 runs per each *K*-value with the highest likelihood estimates. Convergence across runs was assessed by checking the 10 retained replicates per *K*-value provided a similar solution in terms of individual probabilities of assignment to a given ancestral population (*q*-values; Gilbert *et al*. 2012). Then, we used the Greedy algorithm in CLUMPP 1.1.2 to align replicated runs of STRUCTURE for the same *K-*value (Jakobsson & Rosenberg 2007). Following Gilbert *et al*. (2012) and Janes *et al*. (2017), we used two statistics to interpret the number of ancestral populations (*K*) that best describes our data: log probabilities of Pr(X|*K*) (Pritchard *et al*. 2000) and Δ*K* (Evanno *et al*. 2005), both calculated in STRUCTURE HARVESTER 0.6.94 (Earl & vonHoldt 2012).

We reconstructed phylogenetic relationships among the ancestral populations of each species using SNAPP 1.5.2 (Bryant *et al*. 2012), a coalescent-based method for species tree estimation, as implemented in BEAST 2.6.7 (Bouckaert *et al*. 2014). We limited the influence of individuals of admixed ancestry by restricting analyses to conspecific individuals with a high probability of assignment (*q*-value >85%) to the West and East ancestral populations within each species. Accordingly, conspecific individuals from the neighbouring sampling sites MOQ and ZAP (West), and IJU and CTE (East) were grouped, on the basis of STRUCTURE results assuming *K* = 2 for each species. This grouping scheme yielded four genetic groups of similar sample size (*T. canariensis*: West, n = 12, East, n = 10; *T. simplex*: West, n = 9, East, n = 10) and representative of the West and East ancestral populations of each species. Afterwards, the .*usnps* file from IPYRAD was edited and converted into a SNAPP input file, which resulted in a dataset including 5,401 bi-allelic unlinked SNPs shared across demes. Following Noguerales *et al*. (2018), analyses were replicated using different values of the shape (α) and inverse scale (β) parameters of the gamma prior distribution (α = 2, β = 200; α = 2, β = 2,000; α = 2, β = 20,000) for the population size parameter (θ). The forward (*u*) and reverse (*v*) mutation rates were set to be calculated by SNAPP. We used the log-likelihood correction, sampled the coalescent rate and left default settings for all other parameters. We ran two independent runs for each gamma distribution using different starting seeds for 1 million MCMC generations, sampling every 1000 steps (∼1000 genealogies). We used TRACER 1.7 (Rambaut *et al*. 2018) to examine log files, check stationarity and convergence of the chains and confirm that effective sample sizes (ESS) for all parameters were >200. We removed 10% of trees as burn-in and combined tree and log files for replicated runs using LOGCOMBINER 2.4.7 (Drummond & Rambaut 2007). Maximum credibility trees were obtained using TREEANNOTATOR 2.4.7 (Drummond & Rambaut 2007) and the full set of retained trees was displayed with DENSITREE 2.2.6 (Bouckaert 2010). Finally, we used TREESETANALYZER 2.4.7 (Drummond & Rambaut 2007) to estimate the frequency of inferred genealogies contained in the 95% credible set of trees.

#### Methods S4. Testing alternative models of divergence and gene flow

To evaluate the relative statistical support for each of the 9 alternative demographic scenarios (Table S3, Fig. S1), we estimated the composite likelihood of the observed data given a specified model using the site frequency spectrum (SFS) using FASTSIMCOAL 2.5.2.21 (Excoffier *et al*. 2013). We calculated a folded joint SFS using EASYSFS 0.0.1 (I. Overcast, https://github.com/isaacovercast/easySFS). We considered a single SNP per locus to avoid the effects of linkage disequilibrium. These analyses were conducted using the same subset of individuals and demes used for phylogenomic inference in SNAPP (see Supplementary Methods S3).

Each genetic group was downsampled to QR0% of individuals to remove all missing data for the calculation of the SFS, minimise errors with allele frequency estimates, and maximise the number of variable SNPs retained. The final SFS contained 2,730 variable SNPs. Owing to we did not include invariable sites in the SFS, we used the *‘removeZeroSFS’* option in FASTSIMCOAL and fixed the effective population size (*N*_E_) for the West deme of *Tarphius canariensis* (θ_C-W_) to enable the estimation of other parameters (Excoffier *et al*. 2013; Papadopoulou & Knowles 2015; Noguerales & Ortego 2022). We calculated *N*_E_ using nucleotide diversity (π) and estimates of mutation rate per site per generation (μ), where *N*_E_ = π/4μ. We estimated π using phased data from polymorphic and non-polymorphic loci contained in the .*allele* file from IPYRAD using DNASP 6.12.03 (Rozas *et al*. 2017). As for previous analyses, we considered a mutation rate per site per generation of 2.8×10^-9^ (Keightley *et al*. 2014).

Each model was run 100 replicated times considering 100,000-250,000 simulations for the calculation of the composite likelihood, 10-40 expectation-conditional maximisation (ECM) cycles, and a stopping criterion of 0.001 (Excoffier *et al*. 2013). We used an information-theoretic model selection approach based on Akaike’s information criterion (AIC) to determine the probability of each model given the observed data (Burnham & Anderson 2002; *e.g*., Thomé & Carstens 2016). After the composite likelihood was estimated for each model in every replicate, we calculated the AIC scores as detailed in Thomé and Carstens (2016). AIC values for each model were rescaled (AIC) calculating the difference between the AIC value of each model and the minimum AIC obtained among all competing models (*i.e*., the best model has ΔAIC = 0). Point estimates of the different demographic parameters for the best-supported model were selected from the run with the highest maximum composite likelihood. Finally, we calculated confidence intervals (based on the percentile method; *e.g*., de Manuel *et al*. 2016) of parameter estimates from 100 parametric bootstrap replicates by simulating SFS from the maximum composite likelihood estimates and re-estimating parameters each time (Excoffier *et al*. 2013).

#### Methods S5. Analyses of *Wolbachia* infection

Once raw ddRADseq data was demultiplexed in IPYRAD, we searched for bacterial DNA sequences within the .*fastq* file of each individual using CENTRIFUGE 1.0.4 (Kim *et al*. 2016), a microbial classification tool that enables accurate and rapid classification of DNA sequences through applying an indexing scheme based on the Burrows-Wheeler transform and the Ferragina-Manzini index. This novel approach has been proven to provide reliable and sensitive inferences on the presence and abundance of bacterial endosymbionts in ddRADseq data from insect samples (Hinojosa *et al*. 2019, 2020, 2022, 2023). CENTRIFUGE was run with default settings and using the most updated Archaea and Bacteria index provided by CENTRIFUGE developers (https://genome-idx.s3.amazonaws.com/centrifuge/p_compressed_2018_4_15.tar.gz), which is composed of 3,333 complete reference genomes. Raw reads classified as *Wolbachia* by CENTRIFUGE were extracted using MMGBLASTFILTER (T. J. Creedy, https://github.com/tjcreedy/MMGscripts) and quality checked with FASTQC 0.11.7 (Andrews 2010). Then, we searched for and trimmed Illumina adapters using TRIMOMMATIC 0.39 (Bolger *et al*. 2014). After trimming restriction overhangs for enzymes *EcoR1* and *MseI* with CUTADAPT 3.5 (Martin 2011), we used SEQTK 1.3 (H. Li, https://github.com/lh3/seqtk) to trim read ends through applying a conservative error threshold of 0.01, and discard those reads shorter than 50 bp. Resulting quality filtered reads were converted to .*fasta* files using CUTADAPT. Finally, the retained *Wolbachia* sequences were further verified with the *blastn* tool from BLAST+2.15.0 (Camacho *et al*. 2009; *e.g*., Lucek *et al*. 2020) against the NCBI GenBank nucleotide collection (*nt,* 18/02/2024), and only those unambiguously identified as *Wolbachia* were *de novo* assembled, visually inspected and curated using GENEIOUS PRIME 2021.1.1. After assembly, single unmatching *Wolbachia* sequences that were only retrieved in one individual were discarded from downstreaming analyses.

## SUPPLEMENTARY TABLES AND FIGURES

**Table S1.** Geographic coordinates and elevation (metres above sea level) for each of the sampling sites. For each sampling site the number of genotyped individuals per species for ddRADseq and mitochondrial marker is indicated.

| Population code | Sampling site | Longitude | Latitude | Elevation | *T. canariensis (Tca)* |  | *T. simplex (Tsi)* |  |
| --- | --- | --- | --- | --- | --- | --- | --- | --- |
|  |  |  |  |  | N of *Tca*<br>(ddRADseq) | N of *Tca*<br>(mtDNA) | N of *Tsi*<br>(ddRADseq) | N of *Tsi*<br>(mtDNA) |
| MOQ | T19 | -16.308815 | 28.536802 | 778 | 5 | 5 | 2 | 1 |
| ZAP | T18 | -16.296557 | 28.535704 | 883 | 7 | 9 | 7 | 5 |
| AGU | T17 | -16.269782 | 28.533370 | 887 | 7 | 7 | 6 | 3 |
| NCC | T16 | -16.232687 | 28.538624 | 831 | 5 | 5 | 5 | 5 |
| TAG | T15 | -16.226058 | 28.544338 | 808 | 6 | 6 | 5 | 3 |
| ANE | T14 | -16.225111 | 28.540000 | 886 | 5 | 5 | 5 | 2 |
| BQU | T13 | -16.222923 | 28.541069 | 774 | 5 | 5 | 3 | 4 |
| BCA | T12 | -16.219854 | 28.542549 | 739 | 7 | 9 | 6 | 6 |
| FAJ | T11 | -16.215988 | 28.546845 | 701 | 6 | 6 | 7 | 7 |
| GBN | T10 | -16.209503 | 28.547270 | 713 | 5 | 5 | 1 | 3 |
| CEB | T09 | -16.207486 | 28.548379 | 681 | 6 | 7 | - | - |
| EBA | T08 | -16.203886 | 28.549813 | 675 | 7 | 7 | 7 | 7 |
| AMA | T07 | -16.201273 | 28.549903 | 716 | 7 | 7 | 7 | 7 |
| CAG | T06 | -16.196037 | 28.550428 | 707 | 6 | 8 | - | - |
| NPI | T05 | -16.194065 | 28.551440 | 705 | 4 | 5 | - | - |
| PIJ | T04 | -16.189222 | 28.551972 | 791 | 5 | 4 | 5 | 4 |
| CHI | T03 | -16.173594 | 28.559029 | 879 | 5 | 7 | 5 | 3 |
| CTE | T02 | -16.171389 | 28.562028 | 805 | 5 | 5 | 5 | 4 |
| IJU | T01 | -16.169194 | 28.560194 | 755 | 5 | 5 | 5 | 4 |

**Table S2.** Volume and concentration of reagents used for amplifying the COI mitochondrial gene. PCR conditions are also detailed.

| PCR reagents | Concentration | Volume (25 ul) |
| --- | --- | --- |
| Buffer (*MyTaq*<sup>TM</sup>) | 1× | 2.5 ul |
| MgCl<sub>2</sub> (*MyTaq*<sup>TM</sup>) | 3 mM | 1.5 ul |
| dNTPs | 0.4 mM | 1 ul |
| BSA | 0.4 mg/ml | 0.5 ul |
| *Fol-degen-for* | 0.4 mM | 1 ul |
| *Fol-degen-rev* | 0.4 mM | 1 ul |
| Taq (*MyTaq*<sup>TM</sup>) | 0.5 (U/tube) | 0.1 ul |
| DNA extract | - | 2 ul |
| H<sub>2</sub>O | - | 15.4 ul |

| PCR conditions | Temperature | Time |
| --- | --- | --- |
| Initial denaturation | 94°C | 4 min. |
| Denaturation (42 cycles) | 94°C | 30 sec. |
| Annealing (42 cycles) | 46°C | 35 sec. |
| Extension (42 cycles) | 72°C | 45 sec. |
| Final extension | 72°C | 10 min. |

**Table S3.** Comparison of alternative models tested using FASTSIMCOAL (Fig. 4, Fig. S1). Models were constructed assuming no migration (Model A) and migration within species and between species (Models B-C). Interspecific migration was modelled to take place either within western Anaga (W, Models D1-D2), both within western and eastern Anaga (W-E, Models E1-E2) or within and among western and eastern Anaga (full, Models F1-F2). Models assuming asymmetric interspecific migration (Models D2, E2, F2) were also tested. The best-supported model is highlighted in bold.

| | Intraspecific migration | Interspecific migration | Asymmetric migration | lnL | *k* | AIC | ΔAIC | $\omega_i$ |
| --- | --- | --- | --- | --- | --- | --- | --- | --- |
| Model A |  |  |  | -4307.91 | 9 | 8633.82 | 187.99 | 0.00 |
| Model B | ✓ |  |  | -4287.32 | 11 | 8596.64 | 150.81 | 0.00 |
| Model C | ✓ | ✓ |  | -4265.54 | 12 | 8555.09 | 109.26 | 0.00 |
| Model D1 | ✓ | ✓ (W) |  | -4218.61 | 13 | 8463.23 | 17.40 | 0.00 |
| Model D2 | ✓ | ✓ (W) | ✓ | -4217.60 | 15 | 8465.19 | 19.36 | 0.01 |
| **Model E1** | ✓ | ✓ (W-E) |  | **-4208.92** | **14** | **8445.83** | **0.00** | **0.96** |
| Model E2 | ✓ | ✓ (W-E) | ✓ | -4210.77 | 17 | 8455.53 | 9.70 | 0.01 |
| Model F1 | ✓ | ✓ (full) |  | -4210.32 | 16 | 8452.64 | 6.81 | 0.03 |
| Model F2 | ✓ | ✓ (full) | ✓ | -4211.06 | 21 | 8464.13 | 18.29 | 0.00 |

lnL = maximum likelihood estimate of the model; *k* = number of parameters in the model; AIC = Akaike's information criterion value; ΔAIC value from that of the strongest model; $\omega_i$ = AIC weight.

**Table S4.** Results of *Wolbachia* detection in ddRADseq data using CENTRIFUGE and BLAST+. For each species, we provide the number of reads and loci assigned to *Wolbachia* according to CENTRIFUGE and further verified using BLAST+. For the total of *Wolbachia* loci, we provide the number of invariant and variant *Wolbachia* loci sampled across individuals from either one species or both species.

|  | Total | *Tarphius*<br>*canariensis* | *Tarphius*<br>*simplex* |
| --- | --- | --- | --- |
| *Wolbachia* reads (CENTRIFUGE) | 10,614 | 4,286 | 6,328 |
| Verified *Wolbachia* reads (BLAST+) | 10,229 | 4,128 | 6,101 |
| Unverified *Wolbachia* reads (BLAST+) | 385 | 158 | 227 |
| Assembled *Wolbachia* reads (GENEIOUS) | 10,075 | 4,081 | 5,994 |
| Unassembled *Wolbachia* reads (GENEIOUS) | 154 | 47 | 107 |
| *Wolbachia* loci after assembly (GENEIOUS) | 383 | - | - |
| Invariant *Wolbachia* loci | 372 | - | - |
| Invariant *Wolbachia* loci sampled in both species | 125 | - | - |
| Invariant *Wolbachia* loci only sampled in one species | - | 124 | 123 |
| Variant *Wolbachia* loci | 11 | - | - |
| Variant *Wolbachia* loci among individuals within the same species | - | 1 | 3 |
| Variant *Wolbachia* loci among individuals from both species | 7 | - | - |
| Infected individuals after filtering and curation | 25 | 8 | 17 |

**Table S5.** Results of *Wolbachia* detection in ddRADseq data using CENTRIFUGE and BLAST+. For each individual, we provide (a) the number of total host reads in the raw data and mean depth per host locus before and after (in parenthesis) IPYRAD filtering, and (b) the number of reads (*Wb* reads) and loci (*Wb* loci) assigned to *Wolbachia* according to CENTRIFUGE and further verified using BLAST+. For this subset of host individuals with confirmed evidence of *Wolbachia* infection, we provide (c) the number of unique and shared *Wolbachia* loci across individuals.

| Species | Population code | Sampling site | Specimen code | Total raw reads<sup>(a)</sup> | Mean depth per locus<sup>(a)</sup> | *Wb* reads<sup>(b)</sup> | *Wb* loci<sup>(b)</sup> | *Wb* shared loci<sup>(c)</sup> | *Wb* unique loci<sup>(c)</sup> |
| --- | --- | --- | --- | --- | --- | --- | --- | --- | --- |
| *T. canariensis* | MOQ | T19 | caT19MOQ02 | 2081301 | 8.72 (20.35) | 33 | 6 | 2 | 4 |
| *T. canariensis* | MOQ | T19 | caT19MOQ05 | 760772 | 7.13 (15.08) | 1 | 1 | 1 | 0 |
| *T. canariensis* | AGU | T17 | caT17AGU07 | 1286335 | 15.58 (30.36) | 1 | 1 | 1 | 0 |
| *T. canariensis* | TAG | T15 | caT15TAG02 | 2759461 | 13.47 (30.38) | 4083 | 253 | 132 | 121 |
| *T. canariensis* | FAJ | T11 | caT11FAJ01 | 667548 | 6.27 (14.77) | 1 | 1 | 1 | 0 |
| *T. canariensis* | CHI | T03 | caT03CHI05 | 1604555 | 14.18 (27.03) | 1 | 1 | 1 | 0 |
| *T. canariensis* | CTE | T02 | caT02CTE01 | 3514652 | 19.21 (39.93) | 3 | 1 | 1 | 0 |
| *T. canariensis* | IJU | T01 | caT01IJU04 | 1162595 | 7.56 (16.33) | 2 | 1 | 1 | 0 |
| *T. simplex* | MOQ | T19 | siT19MOQ01 | 973201 | 8.82 (17.46) | 1 | 1 | 1 | 0 |
| *T. simplex* | ZAP | T18 | siT18ZAP05 | 3399496 | 15.26 (36.03) | 2 | 2 | 1 | 1 |
| *T. simplex* | ZAP | T18 | siT18ZAP07 | 1503877 | 17.77 (34.90) | 17 | 2 | 0 | 2 |
| *T. simplex* | AGU | T17 | siT17AGU05 | 2324375 | 10.50 (26.52) | 5 | 1 | 1 | 0 |
| *T. simplex* | NCC | T16 | siT16NCC02 | 3901356 | 13.40 (25.39) | 20 | 1 | 1 | 0 |
| *T. simplex* | TAG | T15 | siT15TAG05 | 3145850 | 14.10 (31.99) | 7 | 1 | 1 | 0 |
| *T. simplex* | TAG | T15 | siT15TAG06 | 565065 | 8.37 (15.40) | 413 | 42 | 37 | 5 |
| *T. simplex* | BCA | T12 | siT12BCA01 | 710181 | 4.77 (12.40) | 6 | 1 | 0 | 1 |
| *T. simplex* | BCA | T12 | siT12BCA02 | 1410301 | 15.07 (28.76) | 2812 | 123 | 109 | 14 |
| *T. simplex* | BCA | T12 | siT12BCA04 | 1694588 | 7.84 (20.29) | 2784 | 220 | 142 | 78 |
| *T. simplex* | EBA | T08 | siT08EBA04 | 1610141 | 9.00 (20.94) | 6 | 1 | 1 | 0 |
| *T. simplex* | EBA | T08 | siT08EBA06 | 1007593 | 8.60 (19.42) | 1 | 1 | 1 | 0 |
| *T. simplex* | AMA | T07 | siT07AMA01 | 1605943 | 8.41 (20.18) | 4 | 1 | 1 | 0 |
| *T. simplex* | AMA | T07 | siT07AMA03 | 1693563 | 6.87 (18.60) | 1 | 1 | 1 | 0 |
| *T. simplex* | PIJ | T04 | siT04PIJ01 | 2880694 | 13.49 (28.61) | 3 | 1 | 0 | 1 |
| *T. simplex* | CTE | T02 | siT02CTE05 | 2261906 | 12.55 (24.13) | 16 | 1 | 1 | 0 |
| *T. simplex* | IJU | T01 | siT01IJU02 | 3316659 | 13.96 (33.03) | 1 | 1 | 1 | 0 |

**Figure S1.**
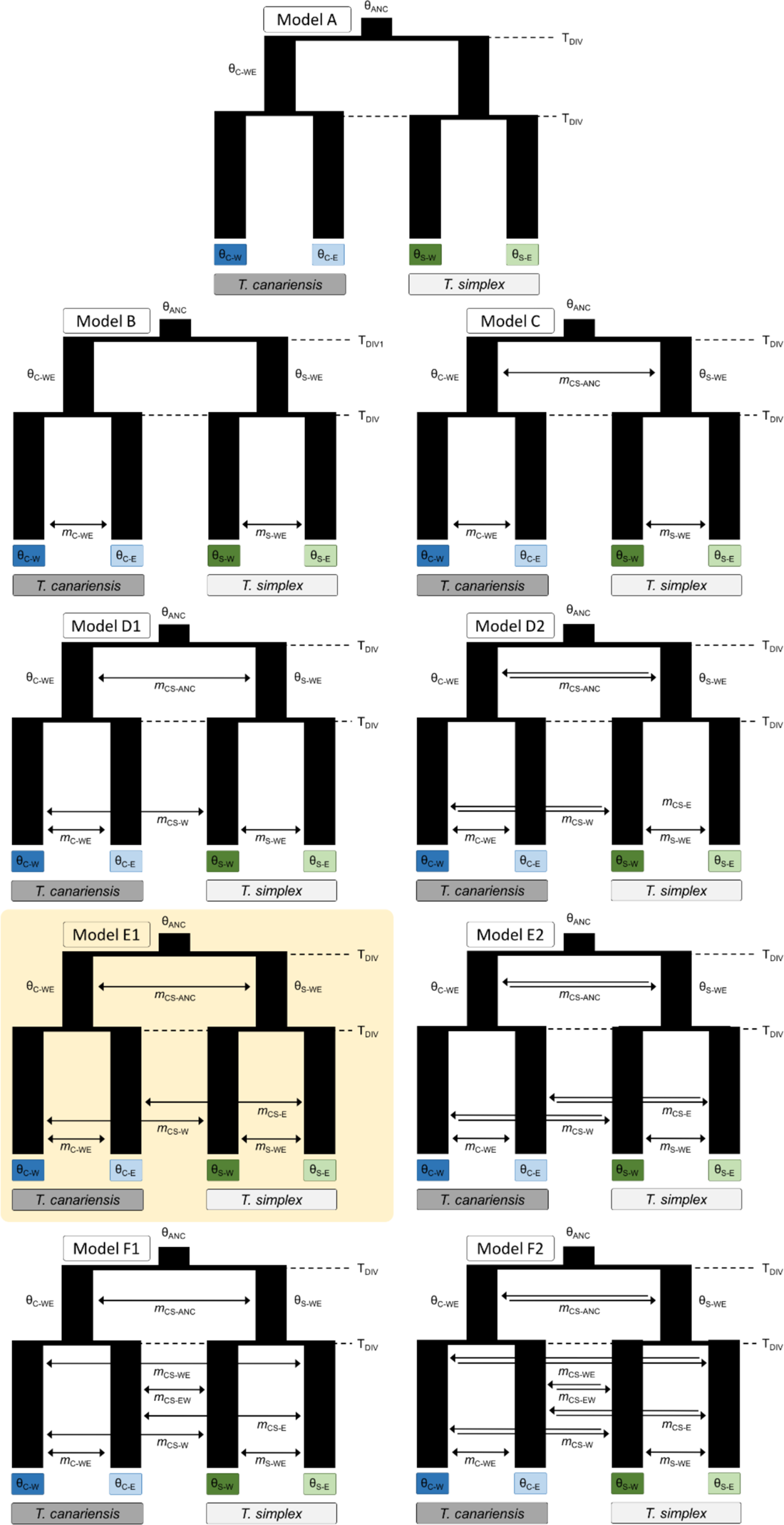
Demographic scenarios tested with FASTSIMCOAL, which were constructed assuming an increasing pattern of migration events within and between species. Model parameters include ancestral and contemporary effective population sizes (θ), divergence times (T_DIV_), and migration rates per generation (*m*). The best-supported model is highlighted.

**Figure S2.**
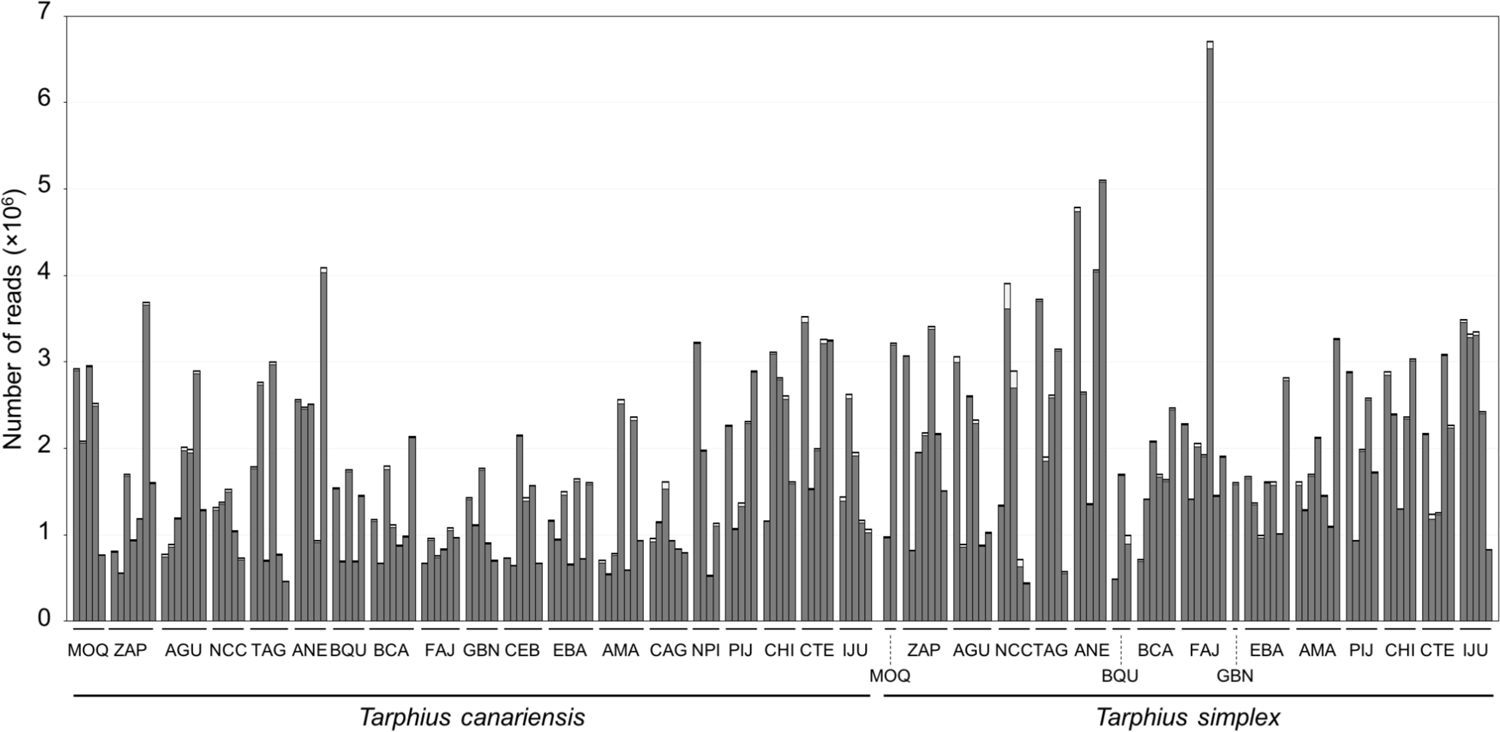
Number of reads per individual before and after different quality filtering steps by IPYRAD. The cumulative stacked bars represent the total number of raw reads obtained for each individual. Within each bar, the pale grey colour represents the reads that were discarded due to short length (*filter_min_trim_len*). Black colour represents a very small proportion of the reads that were subsequently discarded due to not complying with the quality criteria (*max_low_qual_bases*). Finally, the dark grey colour represents the total number of retained reads used to identify homologous loci during the subsequent steps performed in IPYRAD. Population codes as in Table S1.

**Figure S3.**
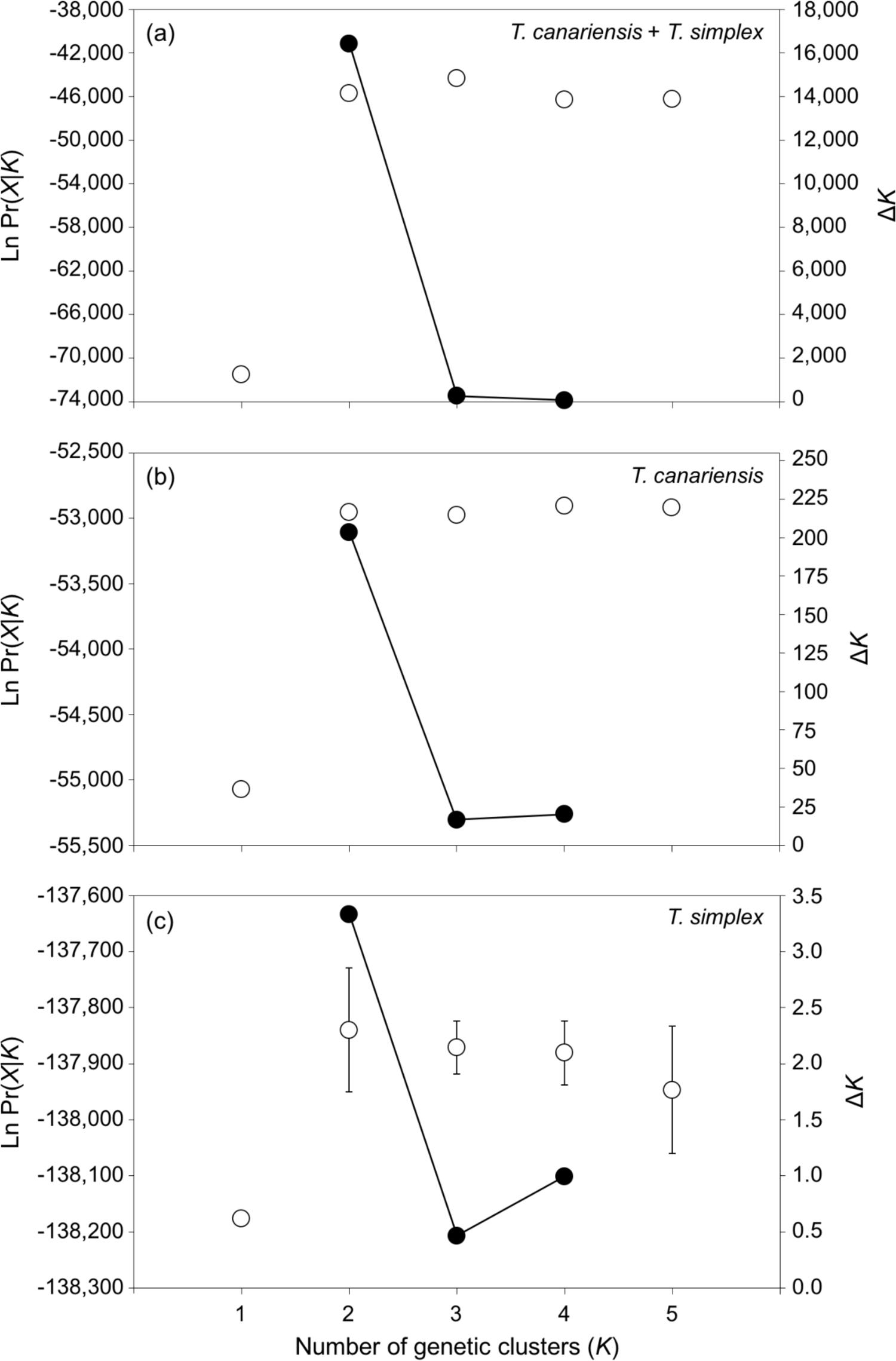
Mean (±SD) log probability of the data (LnPr(X|*K*)) over 10 runs of STRUCTURE (left axes, open dots and error bars) for each value of *K* and the magnitude of Δ*K* (right axes, black dots and continuous line) for analyses including (a) all individuals from the two species *T. canariensis* and *T. simplex*, (b) only individuals of *T. canariensis*, and (c) only individuals of *T. simplex* (Fig. 1).

**Figure S4.**
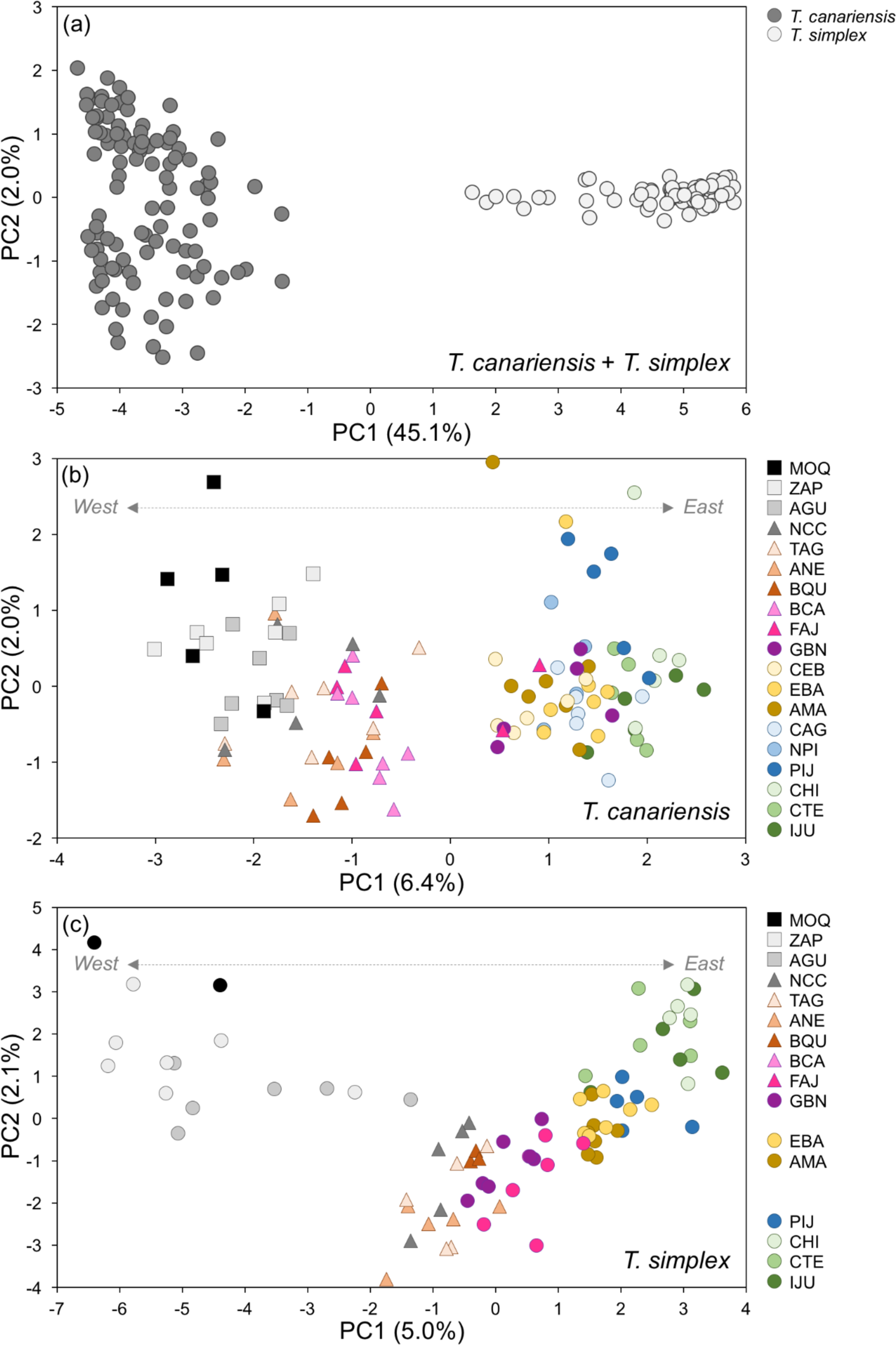
Principal component analysis (PCA) summarising the genetic variation between *T. canariensis* and *T. simplex* (panel a) and within each of the two species (panel b and c, respectively). Population codes as in Table S1.

**Figure S5.**
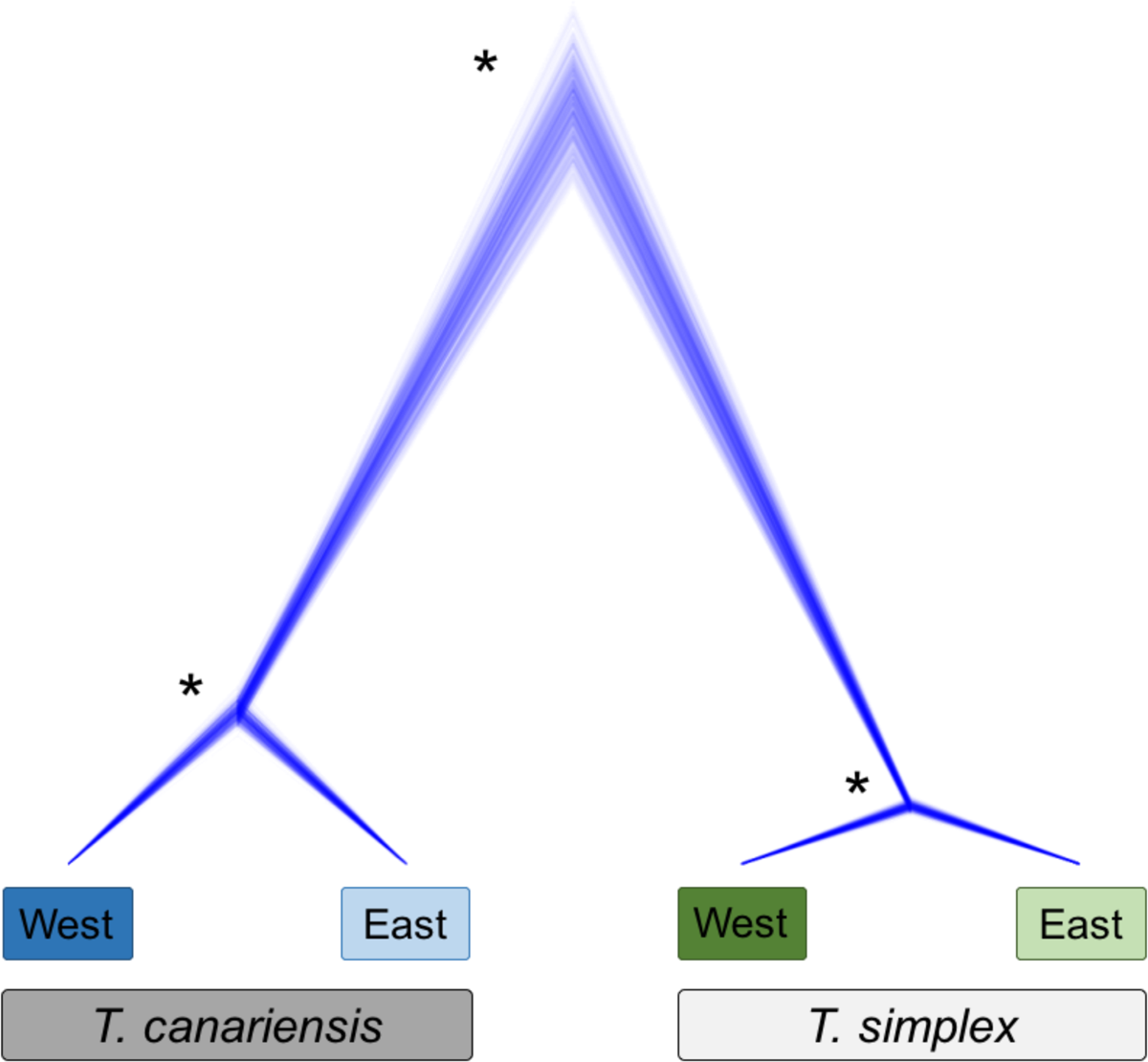
Phylogenetic relationships and branch lengths as inferred in SNAPP among the two main ancestral populations for each of the two species according to STRUCTURE inferences. Asterisks denote fully supported nodes.

**Figure S6.**
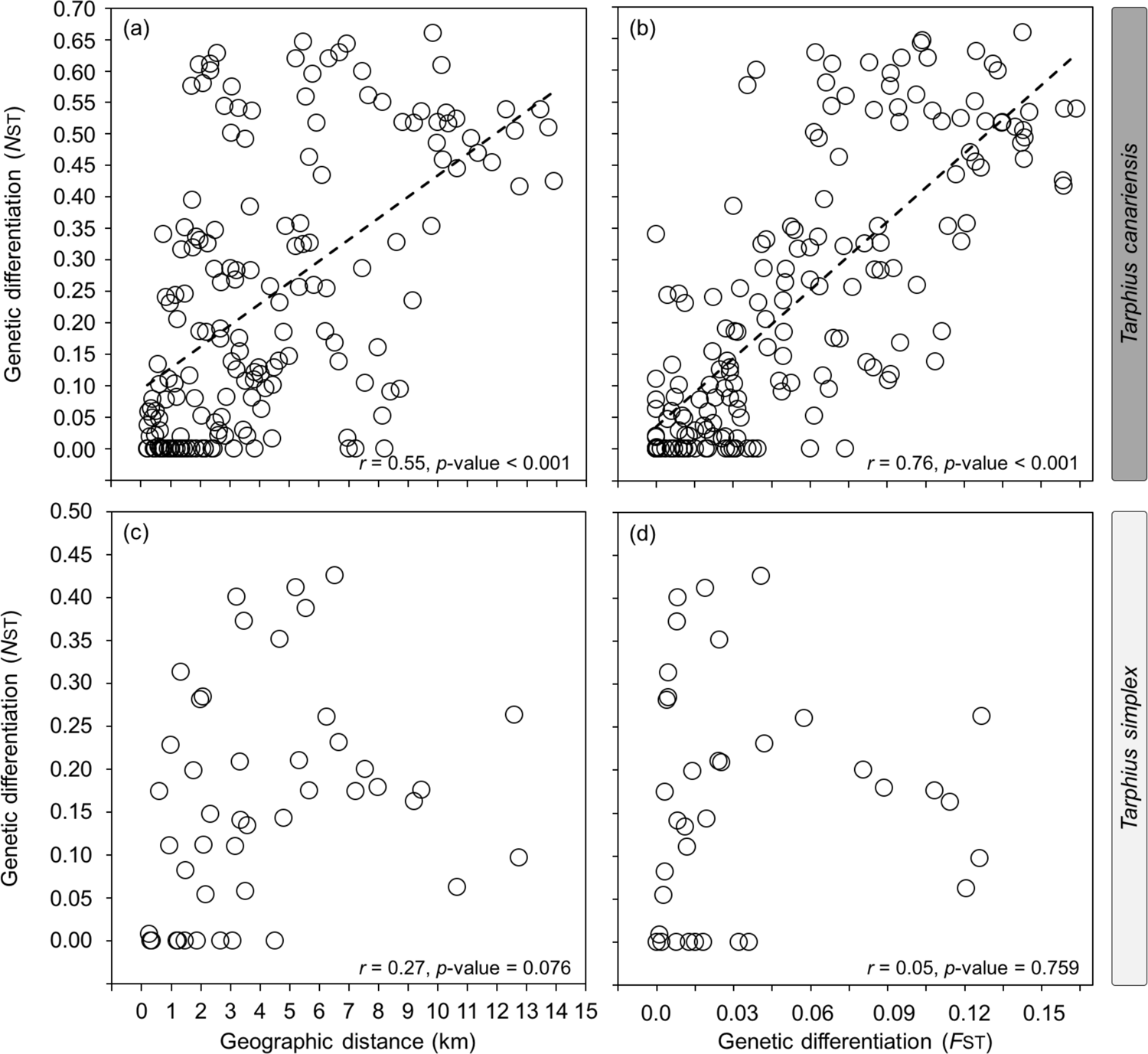
Relationship between genetic differentiation based on mitochondrial data (*N*_ST_), nuclear data (*F*_ST_), and geographic distance between populations of each of the two species, *T. canariensis* (panels a, b) and *T. simplex* (panels c, d).

**Figure S7.**
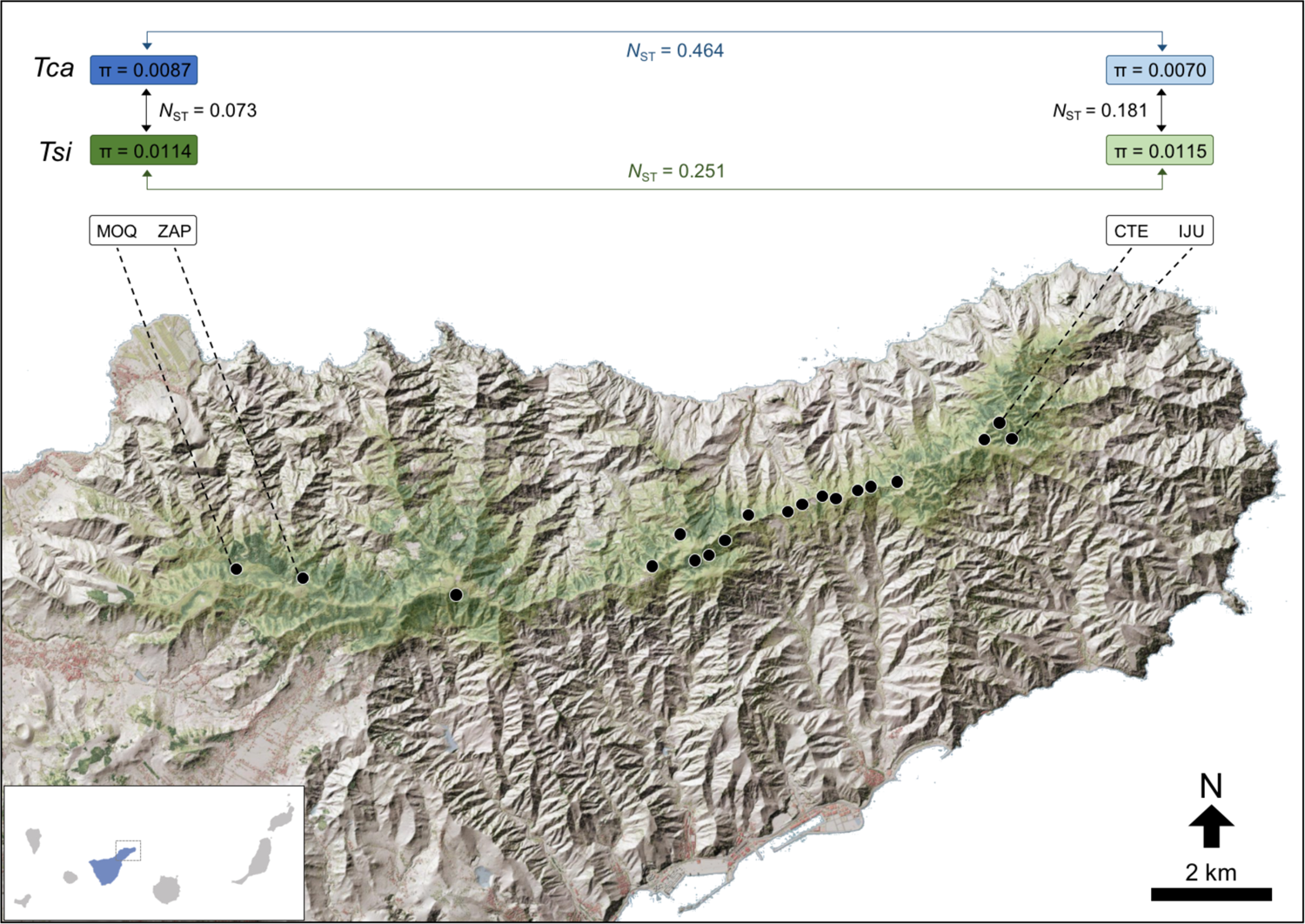
Schematic representation depicting the overall pattern of genetic diversity (π) differentiation (*N*_ST_) within and between species of *Tarphius canariensis* (*Tca*) and *T. simplex* (*Tsi*) using representative populations of its distribution range and mitochondrial data. Population codes as in Table S1.

**Figure S8.**
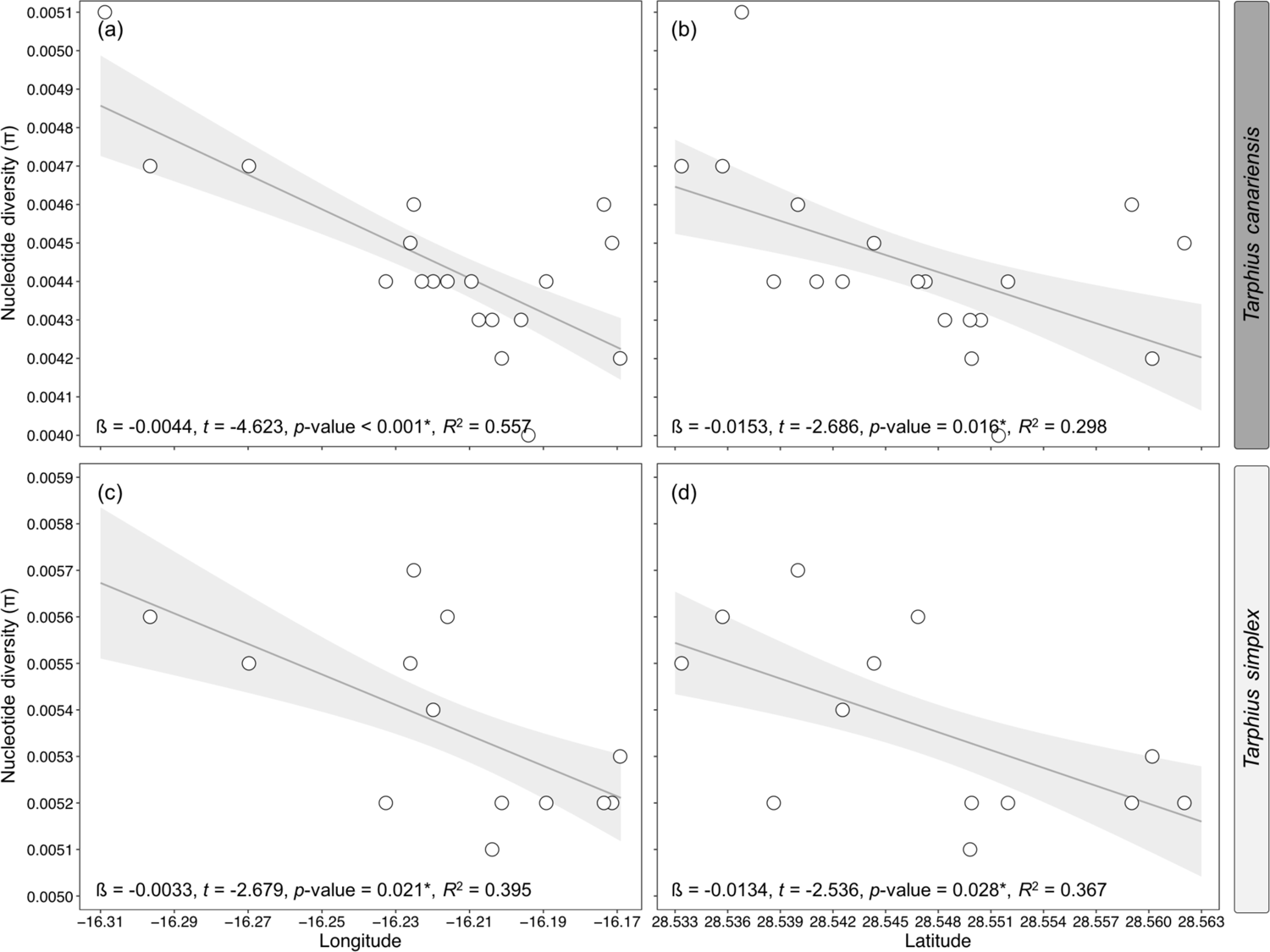
Relationship between nucleotide diversity (π) calculated for the sampling sites of each of the two species, *T. canariensis* (panels a, b) and *T. simplex* (panels c, d), and the spatial variables longitude and latitude. Regression lines and confidence intervals are for significant (*) models. Cohen’ pseudo-*R*^2^ was used to estimate goodness of model fit by calculating [1 - (residual deviance/null deviance)].

**Figure S9.**
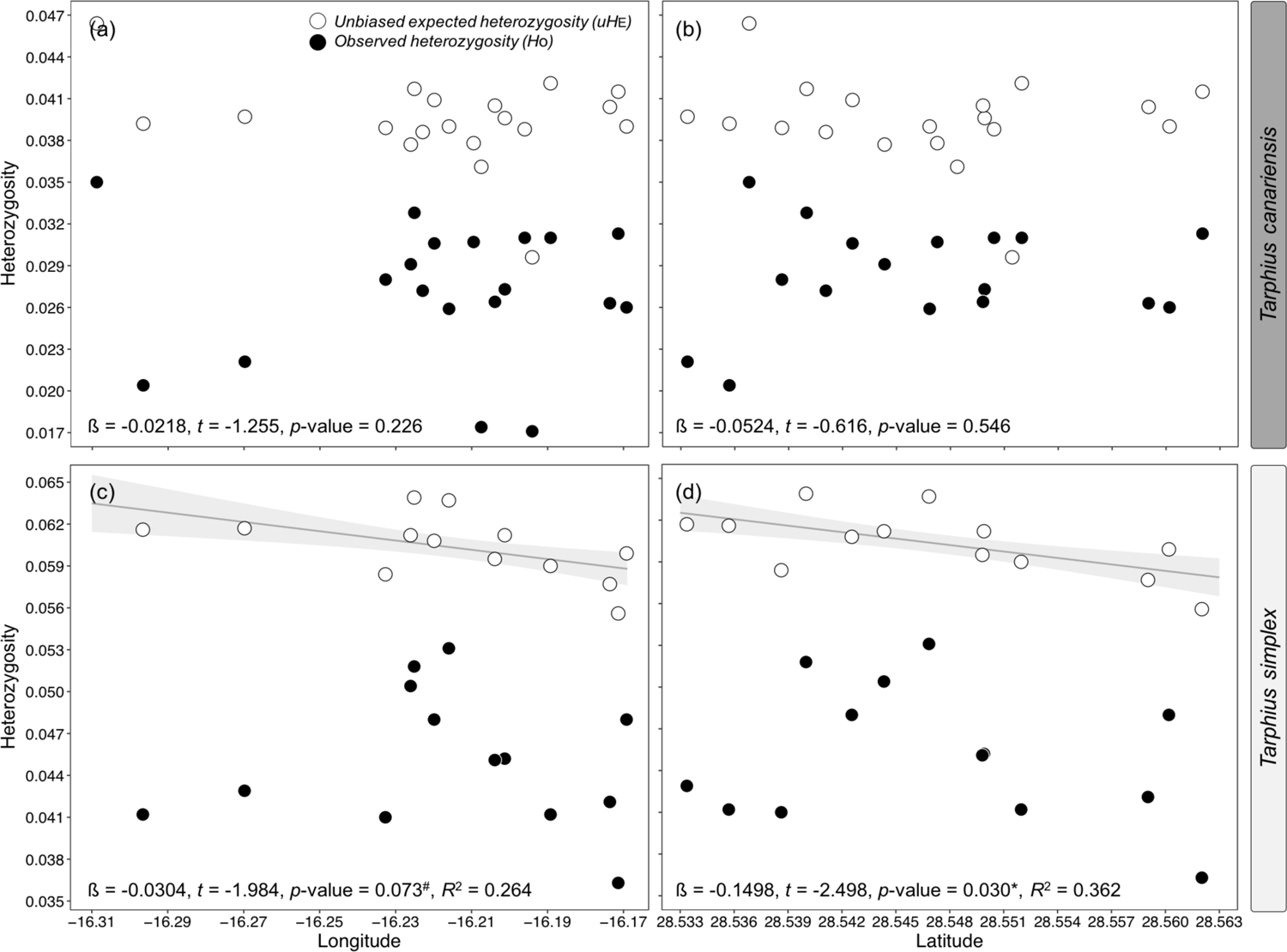
Relationship between unbiased expected heterozygosity (*uH*_E_) calculated for the sampling sites of each of the two species, *T. canariensis* (panels a, b) and *T. simplex* (panels c, d), and the spatial variables longitude and latitude. Regression lines and confidence intervals are only shown for significant (*) and partly significant (#) models. Cohen’ pseudo-*R*^2^ was used to estimate goodness of model fit by calculating [1 - (residual deviance/null deviance)]. Observed heterozygosity (*H*_O_) for each sampling site are shown in black dots.

